# Unraveling Genomic Signatures of lncRNA Expression in Zebrafish Caudal Fin Regeneration: Bridging Regenerative Potential and Positional Memory

**DOI:** 10.1101/2025.11.17.688827

**Authors:** Soumyadeep Paul, A Hariharan, Dasari Abhilash, Surbhi Kohli, Shilpi Minocha, Ishaan Gupta

**Author notes:** first authors.

## Abstract

Zebrafish (Danio rerio) is an essential model organism for tissue regeneration, but the function of long non-coding RNAs (lncRNAs) in tissue regeneration is not well understood. Identifying important regeneration-associated lncRNAs may allow for cross-species applications, improving tissue repair in other organisms. However, few studies and a lack of functional validation complicate our understanding of their mechanistic functions. This research explores novel lncRNAs associated with caudal fin regeneration and positional memory, aiming to identify those that are evolutionarily important across species and may hold universal significance. RNA-seq data deposited in the NCBI database were compared at various important time points (0h post-amputation (hpa), 12 hpa, 1 day post-amputation (dpa), two dpa, three dpa, and seven dpa) and fin parts (proximal, middle, and distal) to uncover major regulatory lncRNAs. Using HISAT2, StringTie, FEELnc, Conservation Analysis and WGCNA (Weighted Gene Co-expression Network Analysis), we found 107 lncRNAs related to regeneration time points and 229 about positional memory during regeneration. Overlapping analysis identified 13 common genomic regions that are complete or partial lncRNAs, indicating a functional connection between regeneration and positional identity and expressed differently in each time point and each position. Additionally, a comparison with regeneration-associated mRNAs revealed that these 13 regions play critical roles in both processes, providing insights into the molecular mechanisms of regenerative precision. Following validation by RT-PCR, the results further suggest that these overlapping regions are differentially expressed across distinct positions within the same tissue, despite a consistent injury pattern. This positional variation in expression may indicate potential roles in both evolutionary adaptation and the regulation of regenerative processes across multiple tissues.

## 1. INTRODUCTION

Long non-coding RNAs (lncRNAs) are RNA molecules greater than 200 nucleotides in length that do not code for proteins, but they help control gene activity and support many important functions inside the cell. In biological systems, lncRNAs act as master regulators of the genome, regulating transcription, post-transcriptional processes, and chromatin remodeling (Ferrer & Dimitrova, 2024). Their role in early embryogenesis, pluripotency induction, and germ cell formation makes them important from a developmental biology context (Llobat, 2021). LncRNAs regulate gene expression at multiple levels, including chromatin modulation, thereby orchestrating embryonic development, cell fate decisions, and complex biological processes, making them key targets for understanding and manipulating development (K. Zhang et al., 2014).

Non-coding RNAs have been recognized as important players in regulating tissue regeneration, coordinating intricate molecular events that indicate regeneration and renewal (Tavares E Silva et al., 2024). LncRNAs play important roles in the different phases of regeneration, from triggering the healing response to directing tissue remodeling, by binding to mRNAs, microRNAs, and proteins (Statello et al., 2021). Their dynamic expression in the various phases of regeneration emphasizes their role in coordinating the regenerative process (Duda et al., 2023). LncRNAs have also been shown to regulate regenerative processes such as cellular senescence, stem cell pluripotency, and differentiation, thus positioning them at the center of the regenerative story (Luo et al., 2016). Recent investigations have identified lncRNAs’ role in transcending cellular senescence, with therapeutic potential in avoiding age-related diseases and enhancing tissue regeneration (Tavares E Silva et al., 2024) . By modulating gene programs directing tissue repair, lncRNAs might represent attractive therapeutic targets to improve healing and age-associated diseases (Zheng et al., 2019).

The identification of universal lncRNAs promoting regeneration across species remains a critical challenge in the field, hindered by their rapid evolution and species-specific functionality (J. Li et al., 2025). The recently developed RegenDbase addresses this by documenting lncRNAs involved in regeneration across organisms like zebrafish, axolotl, and mouse, providing insights into their regulatory control on evolutionarily conserved regeneration pathways (King et al., 2018). However, significant hurdles persist in identifying well-defined regions within the non-coding genome that universally promote or regulate regeneration across species, a limitation that highlights a crucial avenue for future research (J. Li et al., 2025). The complexity of lncRNA functions, coupled with technological and methodological limitations, further complicates this identification process (Mattick et al., 2023). Addressing these challenges requires advancements in technology and methodology, as well as a more comprehensive understanding of lncRNA conservation and function across species (J. Li et al., 2025). The continued development of resources like RegenDbase, combined with cross-species analysis and innovative approaches to evaluate lncRNA conservation, holds promise for unraveling the complex regulatory networks governing tissue regeneration, potentially leading to breakthroughs in regenerative medicine (King et al., 2018).

The ability of human cells to regenerate differs profoundly between organs, with the liver having an outstanding ability to regenerate through hepatocyte cell division and activation of stem cells, while the heart exhibits limited regenerative capacity due to the loss of proliferative capability of cardiomyocytes during early development. This extreme difference between organs in their capacity to regenerate poses challenges and opportunities for the design of site-specific therapeutic approaches to regenerative medicine. In such a scenario, utilizing a model organism to understand pathways and mechanisms involved in regeneration can be quite useful.

Zebrafish have become a prominent model organism for regeneration research due to their remarkable ability to regenerate various tissues, including fins, heart, and brain. It offers unique advantages for studying lncRNA-mediated regeneration (Pansera et al., 2025). This model system has provided valuable insights into the molecular mechanisms underlying tissue repair and regeneration, with studies identifying hundreds of differentially expressed lncRNAs during cardiac and fin regeneration processes (Mathew & Prasad, 2021), offering valuable insights with potential translational applications to human medicine. These fish share approximately 70% of their genes with humans, including 84% of disease-associated genes, making them highly relevant for biomedical research (Howe et al., 2013). Recent studies have revealed hundreds of differentially expressed lncRNAs during zebrafish brain regeneration (Kohli et al., 2024). The remarkable regenerative abilities of zebrafish, coupled with their genetic similarity to humans, have made them invaluable in modeling human diseases and understanding tissue regeneration mechanisms, particularly in areas such as cardiovascular diseases, renal regeneration, and cancer research (Goessling & North, 2014). This research not only elucidates the molecular mechanisms underlying regeneration but also offers promising avenues for developing new therapeutic strategies in human regenerative medicine.

Previous studies have laid a strong foundation for understanding the molecular mechanisms underlying fin regeneration. For instance, research identified 489 genes with differential expression between proximal and distal regenerating tissues, including 29 genes with proximal enrichment and 460 with distal enrichment (Rosati et al., 2024). This spatial regulation of gene expression during regeneration highlights the potential for lncRNAs to play crucial roles in maintaining positional information and guiding the regeneration process. The position-dependent nature of fin regeneration is exemplified by the formation of joints in regenerating fins. The expression of genes such as evx1, dlx5a, and mmp9 is crucial for joint formation and is influenced by the positional context within the fin (Ton & Iovine, 2013). This demonstrates how the location of cells within the fin can significantly affect the regeneration process and outcome.

Furthermore, the study of chromatin modifications during fin regeneration has revealed that H3K4me3 levels increase over gene promoters that become transcriptionally active, recapitulating patterns observed during embryonic development (Macrae et al., 2023). This suggests that regeneration may involve the reactivation of developmental programs, which could be regulated by lncRNAs in a position-specific manner. The study of lncRNAs in zebrafish fin regeneration, with its distinct positional memory and spatially regulated gene expression, offers a unique opportunity to unravel the complex regulatory networks governing tissue regeneration.

Firstly, the following study makes use of the best available transcriptome-wide approaches in identifying and characterizing novel lncRNAs that participate in zebrafish caudal fin regeneration. Integrating RNA-seq data analysis using powerful computational tools with the validation through an experimental approach may allow for providing a complete map of the time-course landscape of lncRNA expression. Our focus is on key time points corresponding to well-defined stages of regeneration, namely the wound healing phase (0-12 hours post-amputation), blastema formation phase (1-2 days post-amputation), and finally, the regenerative outgrowth phase (3-7 days post-amputation).

Secondly, this study aims to further our understanding of the molecular basis of positional memory in zebrafish caudal fin regeneration by analyzing RNA sequencing data to identify and characterize novel lncRNAs associated with spatial identity. By employing advanced bioinformatics tools and statistical analyses, we seek to uncover the distinct expression patterns of lncRNAs along the proximodistal axis and their potential roles in guiding the regeneration process. Our findings may provide new avenues for therapeutic interventions in regenerative medicine, potentially leading to improved strategies for tissue repair and regeneration in humans (Khan et al., 2025) (Bhatt et al., 2025).

At the end, provide a strong perspective on lncRNAs that are crucial for both identifying cell position and playing a significant role in regeneration.

## 2. METHODS

The methodology for this study comprises five primary steps, as depicted in Figure 1 under the heading “Basic Approach Methodology.” Furthermore, Figure 1 also provides a comprehensive schematic of the lncRNA discovery pipeline, detailing each stage involved in the identification and characterization of long non-coding RNAs.

**Figure 1:**
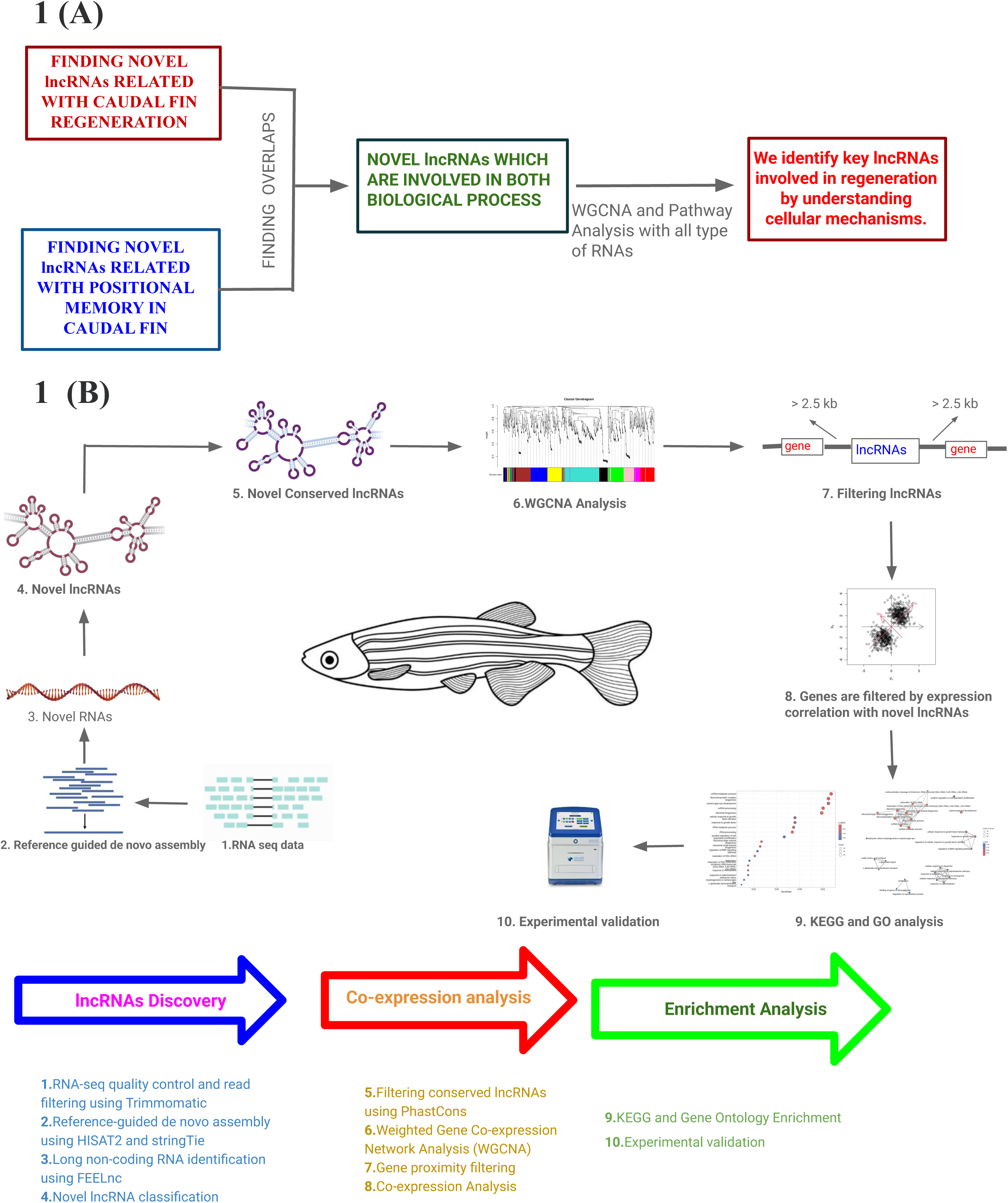
(A) Our Basic Approach Methodology, (B)Schematic of the lncRNA discovery pipeline

### 2.1. NOVEL LNCRNA IDENTIFICATION METHODOLOGY

#### 2.1.1 Data acquisition and sample description

**FOR CAUDAL FIN REGENERATION:**

RNA sequencing data were obtained from the NCBI database under the BioProject accession number PRJNA248169, as part of a study conducted by Banu, S. et al., 2022, (Banu et al., 2022). The data were analyzed to assess lncRNA expression at various time points: 0 hours post-amputation (hpa), 12 hpa, 1-day post-amputation (dpa), 2 dpa, 3 dpa, and 7 dpa. These time points capture the sequential phases of regeneration, beginning with inflammation and wound healing (0–12 hpa), followed by blastema formation and proliferative expansion (1–3 dpa), and culminating in tissue patterning and remodeling (7 dpa). Morphological landmarks were also noted, with bony ray (lepidotrichia) formation evident between 2–3 dpa, and actinotrichia aligned along the proximal–distal axis by 7 dpa (Fig. 2 (A)),(Banu et al., 2022).

**Figure 2.**
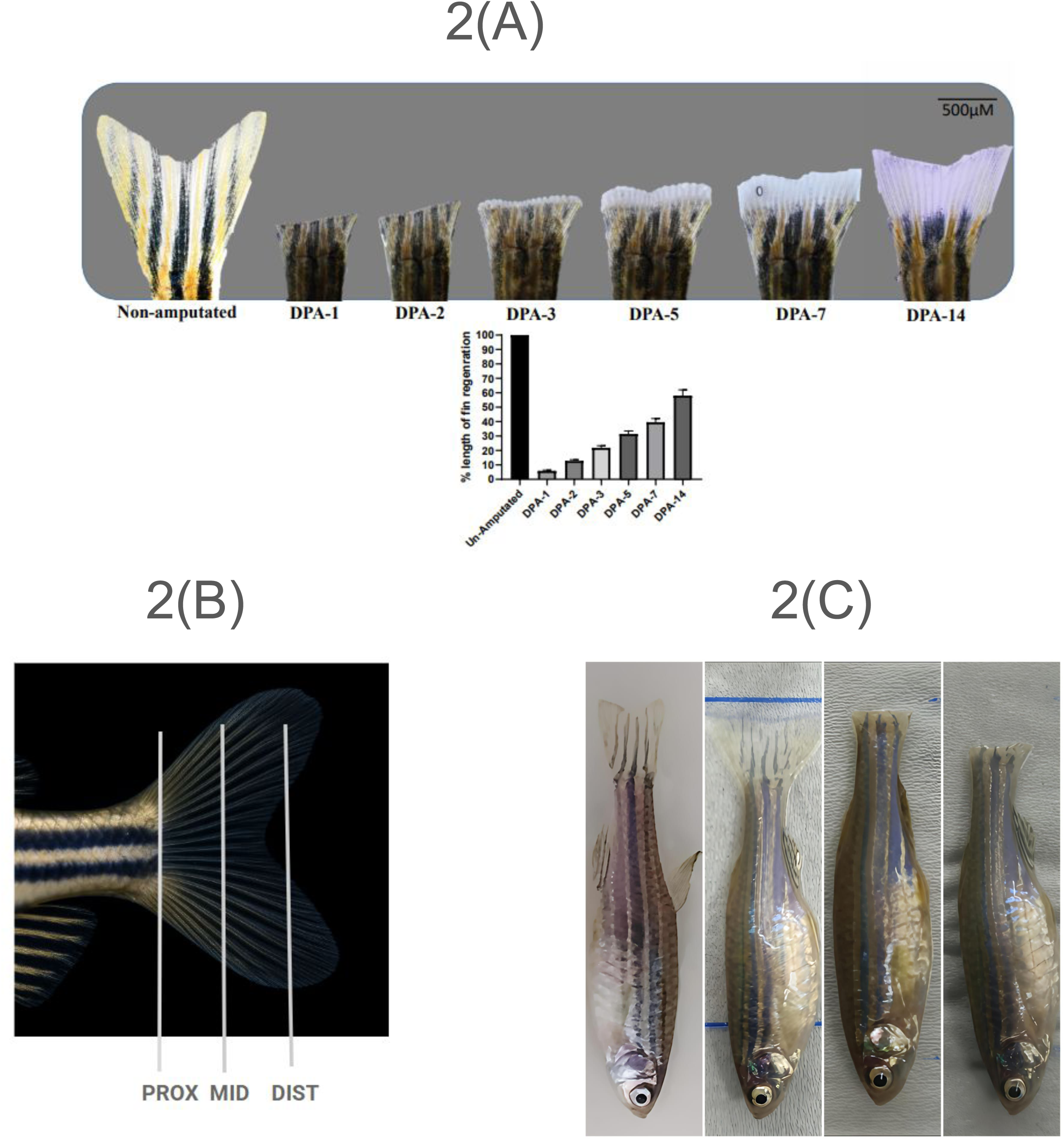
(A) Representative photos illustrating gradual caudal fin regrowth in adult zebrafish after proximal amputation. Caudal fins were severed at the proximal level (0 DPA), and regeneration was observed at consecutive time points: 1, 2, 3, 5, 7, and 14 days post-amputation (DPA). The un-amputated fin (far left) illustrates the baseline full-length fin. Images indicate the steady and ordered re-growth of fin tissue over time, with considerable morphological repair visible by DPA-14. All photos are scaled to the same magnification; scale bar = 500 µm. (B) Schematic representation of the zebrafish caudal fin illustrating the three defined regions along the proximodistal axis: proximal (PROX), middle (MID), and distal (DIST). (C) Representative dorsal view images of adult zebrafish showing: intact caudal fin (first panel), fin amputated at the distal region (second panel), fin amputated at the middle region (third panel), and fin amputated at the proximal region (fourth panel).

**FOR POSITIONAL MEMORY :**

RNA sequencing data utilized in this study were obtained from the Gene Expression Omnibus (GEO) database (accession no. GSE92760) and were originally generated as part of the study, Transcriptomic, Proteomic, and Metabolomic Landscape of Positional Memory in the Caudal Fin of Zebrafish by Rabinowitz et al (Akimenko et al., 1995) (Rabinowitz et al., 2017). The dataset consists of RNA-seq profiles from five biological replicates for each of three distinct regions along the proximodistal axis of the zebrafish caudal fin: proximal, middle, and distal (totaling 15 samples). Each biological replicate represents pooled fin tissue collected from two male and two female zebrafish (Akimenko et al., 1995) (Rabinowitz et al., 2017).

Together, these two datasets enable parallel investigation of temporal lncRNA dynamics during regeneration and spatial lncRNA signatures underlying positional memory, offering a comprehensive view of molecular mechanisms driving zebrafish caudal fin regrowth, which in Figure 2.

#### 2.1.2 Transcriptome Assembly and Mapping

FASTQ files corresponding to each sample were retrieved from the NCBI Sequence Read Archive (SRA) using their respective BioProject accession numbers. The raw sequencing reads underwent quality control and preprocessing using Trimmomatic (v0.38) (Bolger et al., 2014), where adapter sequences were removed and low-quality bases were trimmed using default parameters to ensure high-quality downstream analysis. The resulting cleaned reads were then aligned to the Danio rerio reference genome (GRCz11) using HISAT2 (v2.2.1)(Kim et al., 2015).

During alignment, the --no-templatelen-adjustment parameter was used to disable template length correction, which is more suitable for RNA-seq data and helps maintain accuracy in transcript-level alignment. StringTie (v2.2.1) (Pertea et al., 2015) was used for reference-guided transcript assembly with reverse-stranded libraries, using HISAT2 outputs and merging all sample transcriptomes.

#### 2.1.3 Identification and Classification of lncRNAs

For annotation and classification, we applied FEELnc (v0.2.1) (Wucher et al., 2017), a machine learning–based framework specifically designed to distinguish lncRNAs from protein-coding transcripts. Unlike alignment-dependent methods, FEELnc relies on a Random Forest classifier trained on multi-k-mer frequency patterns and relaxed ORF features, allowing it to capture subtle differences between coding and noncoding sequences. To improve the reliability of our dataset, we used the filtering modules provided within the pipeline: FEELnc_filter.pl to exclude likely artifacts such as mono-exonic transcripts, and FEELnc_codpot.pl to evaluate and remove sequences with coding potential (Wucher et al., 2017). The remaining transcripts were then organized according to their genomic context and proximity to annotated genes, generating a curated set of high-confidence lncRNAs for downstream analysis.

#### 2.1.4 Conservation Analysis

To evaluate whether the lncRNAs we identified showed evidence of evolutionary preservation, we analyzed their sequences with PhastCons (v1.6) (Siepel & Haussler, 2004), which applies a phylogenetic hidden Markov model to estimate conservation scores across species. Because multiple sequence alignment (MSA) data were not directly available for the updated zebrafish genome build, we first converted transcript coordinates to the appropriate reference version using BedFile Liftover, ensuring compatibility with the conservation framework. This allowed us to reliably assess which lncRNAs carry signatures of cross-species conservation, a feature often associated with functional relevance.

#### 2.1.5 Weighted Gene Co-expression Network Analysis (WGCNA)

To explore higher-order relationships among transcripts, we applied Weighted Gene Co-expression Network Analysis (WGCNA) (Langfelder & Horvath, 2008) using the R package (v1.72-5). Prior to network construction, expression values were TPM-normalized to reduce technical bias and make samples comparable. In building the network, we selected a soft-thresholding power of 30, as this value best satisfied the scale-free topology criterion, ensuring that the resulting network captured biologically realistic patterns of connectivity. We focused on a signed network, which gives greater weight to positive correlations, and organized the data with parameters tuned for sensitivity: a deep split level of 2, a minimum module size of 30 transcripts, and a block size capped at 4000 genes. Modules were merged when their similarity exceeded a cut height of 0.25, preventing fragmentation of closely related clusters. Through this process, we obtained groups of co-expressed genes and lncRNAs that highlight coordinated biological activity and point toward regulatory programs relevant to the processes under study.

#### 2.1.6 Gene Proximity Analysis

After identifying key modules through WGCNA, we refined the set of candidate lncRNAs by considering their genomic neighborhood. Using coordinate information extracted from reference GTF annotations and the StringTie assemblies, we checked whether each lncRNA lay within 2.5 kb of a known coding gene. Those falling inside this window were removed from further consideration, as they might represent transcriptional noise or extensions of nearby transcripts rather than independent loci. To further strengthen the functional relevance of the modules, we re-examined patterns of co-expression between lncRNAs and protein-coding genes. Our guiding principle was that transcripts that rise and fall together are more likely to participate in related processes. For this reason, we applied Pearson correlation analysis and retained only strong relationships (correlation coefficient ≥ 0.75). Within these networks, particular emphasis was placed on lncRNAs that repeatedly occupied central or hub-like positions, as these were more likely to play important biological roles.

#### 2.1.7 Pathway Enrichment Analysis

To understand the functional context of the novel lncRNAs uncovered in our dataset, we examined the genes that were consistently co-expressed with them and asked what kinds of biological activities these associations might imply. For this purpose, we relied on the ClusterProfiler (Yu et al., 2012) package in R, which allowed us to link expression clusters to well-annotated functional categories. Instead of simply listing the genes, we traced their connections to established Gene Ontology processes and KEGG signaling pathways, using all expressed transcripts as a reference background to avoid bias. Only associations that passed a conservative statistical cutoff (*p* < 0.05) were considered meaningful. By taking this approach, we were able to highlight functional patterns pointing to roles for the lncRNAs in developmental regulation, tissue physiology, and regenerative biology, while also prioritizing specific candidates for further experimental work.

### 2.2. FINDING OVERLAPS

We aim to identify the overlap between novel lncRNAs that play a crucial role in different biological processes, particularly those involved in fin regeneration and positional memory. By analyzing these lncRNAs, we seek to determine which ones are consistently essential across multiple regenerative stages and spatial identity regulation. This approach allows us to uncover key regulatory lncRNAs that not only drive the regeneration process but also contribute to maintaining positional cues within the tissue. Identifying such lncRNAs could provide deeper insights into the molecular mechanisms underlying regeneration and spatial patterning, potentially leading to broader applications in regenerative medicine.

### 2.3. WGCNA AND PATHWAY ANALYSIS

We performed WGCNA to analyze all RNAs from the RNA-seq dataset on regeneration and positional memory that come from both the NCBI database (PRJNA248169 and GEO accession no. GSE92760), identifying those overlapped lncRNAs that are really crucial for both biological processes. WGCNA (v1.72-5) was used to detect co-expressed genes and lncRNA clusters, with normalized data and a soft-thresholding power of 30, ensuring a scale-free network. A signed network was constructed with a deep split level of 2, a minimum module size of 30 genes, and a maximum block size of 4000 genes, merging related modules at a cut height of 0.25.

Pathway enrichment analysis using ClusterProfiler linked novel lncRNA clusters to key biological functions and pathways. Gene Ontology (GO) and KEGG enrichment were performed with a P-value threshold of <0.05, using all expressed genes as the background. This analysis provided valuable insights into the functional roles of lncRNAs in zebrafish development, regeneration, and positional memory.

### 2.4. Zebrafish maintenance and induction of Stab wound injury

Adult zebrafish (3 months old, 25–30 mm length) were maintained under standard laboratory conditions. For fin amputation, fish were anesthetized in freshly prepared 0.02% Tricaine solution (diluted in system water), with complete anesthesia confirmed when the animals became immobile within 40–60 second (Uemoto et al., 2020) (Srivastava et al., 2024). At 0 days post-amputation (DPA), the caudal fin was amputated at the proximal position, i.e., at the level where the fin rays emerge from the peduncle, using a sterile surgical blade (No. 11 scalpel). All amputations were performed uniformly across experimental animals. Following surgery, fish were transferred to fresh tank water under normal housing conditions to facilitate regeneration. Regenerating fins were imaged at 1, 2, 3, 5, and 7 dpa using a stereo microscope (SMZ1270) equipped with a digital camera. Fin regeneration was quantified using ImageJ software (NIH, USA) by calculating the total regenerated fin area at each time point relative to the pre-amputation fin area (set at 100%).

### 2.5. RNA isolation, cDNA synthesis, and qRT-PCR

The experiments were conducted using AB strain zebrafish (3–4 months old). Total RNA was extracted from tissue samples using Trizol reagent (Invitrogen, 15596026), followed by phase separation with chloroform (Merck, 1.94506.0521) and precipitation with isopropanol (Sigma, I9516). RNA quantity and purity were assessed using a Nanodrop™ One/OneC Microvolume UV-Vis Spectrophotometer (Thermo Scientific, ND-ONE-W). First-strand cDNA was synthesized using the TAKARA cDNA synthesis kit (6110A), and qRT-PCR reactions were set up with iTaq Universal SYBR Green Supermix (Bio-Rad, 1725121). The reactions were performed in hard-shell PCR plates (Bio-Rad, HSP9601) sealed with Bio-Rad plate sealers (MSB1000) on a CFX96 Touch Deep Real-Time PCR Detection System (Bio-Rad, 1854096). Nuclease-free water (Qiagen, 129115) was used for all reactions, and gene-specific real-time primers (Eurofins) were employed for amplification. Tissue homogenization was performed using a Jaisbo MT-13K homogenizer, and sample dissection and phenotypic observation were carried out using a Nikon Trinocular Stereo Zoom Microscope (SMZ1270).

To assess the temporal expression profile of selected long non-coding RNAs (lncRNAs) during caudal fin regeneration, total RNA was isolated from the proximal region of regenerating caudal fins at multiple time points: 1, 2, 3, 5, and 7-day post-amputation (DPA) and for distal and median region we done for time point: 1, 3, and 7 DPA. Fin samples were collected in RNase-free conditions, and RNA extraction was performed using the Trizol (Invitrogen, catalog number: 15596026) according to the manufacturer’s protocol. The purity and concentration of RNA were measured using a NanoDrop™ spectrophotometer. For cDNA synthesis, 1 µg of total RNA from each sample was reverse-transcribed using the TAKARA cDNA synthesis kit (catalog number:6110A) following the recommended protocol. Quantitative real-time PCR (qPCR) was carried out using iTaq Universal SYBR Green Supermix (Biorad, catalog number: 1725121). Each reaction was performed in triplicate. Expression levels of eight selected lncRNAs were quantified, and relative gene expression was calculated using the 2^−ΔΔCt method, with RPL13a as the Internal control.

## 3. RESULTS

### 3.1. Genome-wide discovery and analysis of novel lncRNAs in zebrafish caudal fin regeneration

The RNA sequencing data, sourced from the NCBI database, consisted of 6 samples collected at various time points following caudal fin amputation in zebrafish. These time points included 0 hours post-amputation (hpa), 12 hpa, 1 day post-amputation (dpa), 2 dpa, 3 dpa, and 7 dpa, corresponding to different stages of the regeneration process. Transcriptome-wide profiling enabled the systematic discovery of novel lncRNAs expressed during regeneration. Dynamic expression patterns were observed, reflecting stage-specific regulation of lncRNA networks. Early time points were enriched for lncRNAs associated with immune and wound responses, whereas intermediate and later stages showed induction of lncRNAs linked to blastema proliferation, extracellular matrix organization, and skeletal patterning. This genome-wide catalog of regeneration-associated lncRNAs establishes a framework for exploring their putative roles in coordinating the molecular transitions from wound closure to blastema-driven regrowth and tissue remodeling.

**Transcriptome Assembly and Mapping:**

The initial quality assessment of our raw sequencing data, performed using FastQC, revealed high-quality metrics across all samples, with an average Phred score of more than 30. After trimming adapters and low-quality bases using Trimmomatic, the cleaned data were aligned to the zebrafish reference genome (GRCz11) using HISAT2. This alignment achieved a high mapping efficiency, with an average of 92% of reads successfully aligned. Transcriptome assembly was carried out using StringTie, which integrated the alignments and identified approximately 892,910 transcripts, including both known and novel isoforms.

**Identification of potential LncRNA candidates:**

Utilizing the FEELnc pipeline, we classified transcripts from our de novo transcriptome assembly into coding (mRNAs) and long non-coding RNAs (lncRNAs). FEELnc identified 10,113 putative lncRNAs based on coding potential, multi-k-mer profiles and constrained open reading frame (ORF) characteristics. To investigate their evolutionary trajectory, we employed PhastCons—a phylogenetic hidden Markov model that integrates multiple vertebrate genomes—to quantify conservation. Since lncRNAs exhibit low conservation across different species, it was assumed that lncRNAs with any degree of conservation may harbor pivotal biological functions. Therefore, we specifically focused on the 6,114 lncRNAs that manifested non-zero conservation. We further refined this subset to those with at least 50% overlap within conserved regions and ultimately isolated the top 5% based on conservation metrics Supplementary Document 1. This approach yielded 1,514 lncRNAs, having pronounced conservation signatures. These highly conserved lncRNAs, therefore, could emerge as prime candidates for downstream functional characterization, as their evolutionary preservation suggests potential biological relevance.

We assessed the expression profiles of identified lncRNA candidates across various samples using principal component analysis (PCA) of normalized lncRNA counts (Figure 3). The PCA results showed that samples from 1-day post-lesion (1dpl) distinctly clustered apart from other stages, which were closer to 12hpa. This shows that the novel lncRNAs are highly active immediately after injury and slowly diminish during recovery. This pattern was corroborated by a sample-to-sample distance heatmap, where lower correlations (blue) are seen between 72 hrs and other time points, suggesting a divergence in expression patterns at this stage. These observations suggest that the expression of these lncRNAs is particularly altered during the initial stage of wound healing, possibly playing critical roles in the initial stage of the brain regeneration process, and then regeneration continues. Similar pattern of lncRNAs being active during early injury, while their pattern reduces during late stages in traumatic brain injury in zebrafish (Kohli et al., 2024).

**Figure 3.**
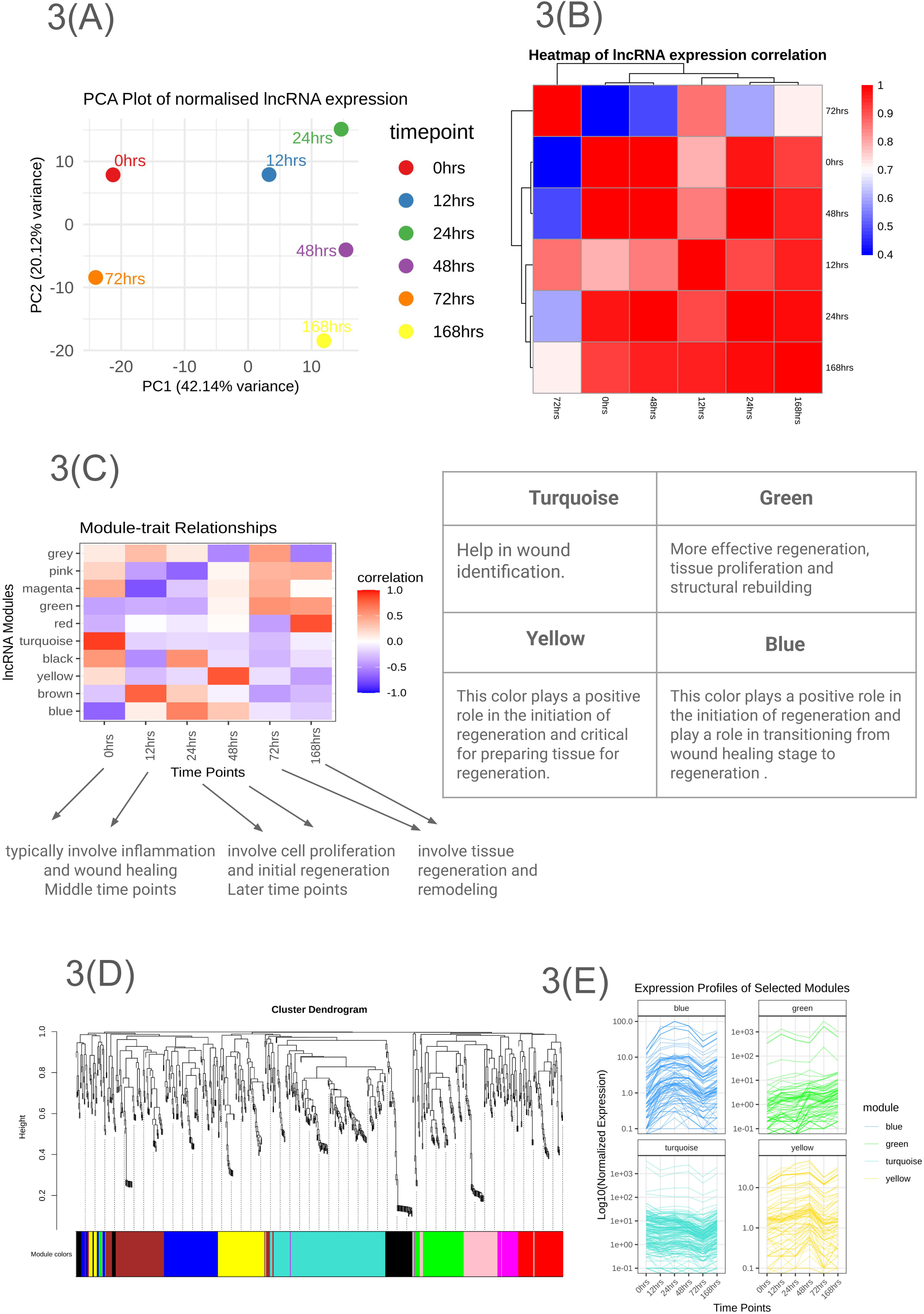
(A) PCA plot of normalized lncRNA expression. (B) Heatmap of lncRNA expression correlation between samples. This heatmap provides insights into how lncRNA expression changes over time, highlighting periods of similarity and divergence in expression profiles. (C) Heatmap showing the correlation of expression profiles among lncRNA clusters. Turquoise, Green, Yellow, and Blue modules show some significant changes. (D) Dendrogram of WGCNA Clusters with 15,14 novel lncRNA. (E) Expression profiles of the individual long non-coding RNAs in the selected modules co-expressed cluster. The Y-axis indicates normalized gene expression, and the X-axis indicates samples labeled according to the treatment.

We performed Weighted Gene Co-expression Network Analysis (WGCNA) to pinpoint clusters of highly coexpressed novel lncRNAs with distinct sample-specific expression patterns (Figure 3). This approach aimed to identify novel lncRNAs that may regulate fin regeneration. Our analysis identified 10 modules with unique expression profiles across the fin injury recovery timeline.

We identified 183 novel lncRNAs in the turquoise module, 79 in the green module, 102 in the blue module, and 85 in the yellow module. To ensure the accuracy of our functional inference, we implemented stringent criteria, excluding any novel lncRNAs located within 2.5 kb of coding genes to avoid potential interference with genes. This filtration left us with 46 lncRNAs in the turquoise module, 20 in the green module, 23 in the blue module, and 18 in the yellow module.

**Table 1:** Gene proximity analysis filtering table.

| MODULE | Gene before proximity Filtering | Gene after proximity Filtering | Transcript before proximity Filtering | Transcript after Proximity Filtering |
| --- | --- | --- | --- | --- |
| TURQUOISE | 120 | 33 | 183 | 46 |
| GREEN | 61 | 16 | 79 | 20 |
| BLUE | 77 | 17 | 102 | 23 |
| YELLOW | 56 | 13 | 85 | 18 |

**Functional significance of identified lncRNAs:**

To further elucidate the biological relevance of these newly identified lncRNAs, we extended our analysis by correlating their expression profiles with those of established protein-coding genes. We prioritized genes exhibiting a robust correlation (|r| > 0.9) with the lncRNAs and subsequently performed Gene Ontology (GO) and KEGG pathway enrichment analyses. This approach aimed to uncover the biological processes and pathways potentially modulated by these lncRNAs during the early wound-healing phase of brain regeneration. Notably, genes coexpressed with lncRNAs in the brown module were enriched for metabolic pathways pertinent to RNA splicing and peptide biosynthetic processes. This suggests that these lncRNAs may be pivotal in orchestrating these foundational cellular activities (Figure 4).

**Figure 4.**
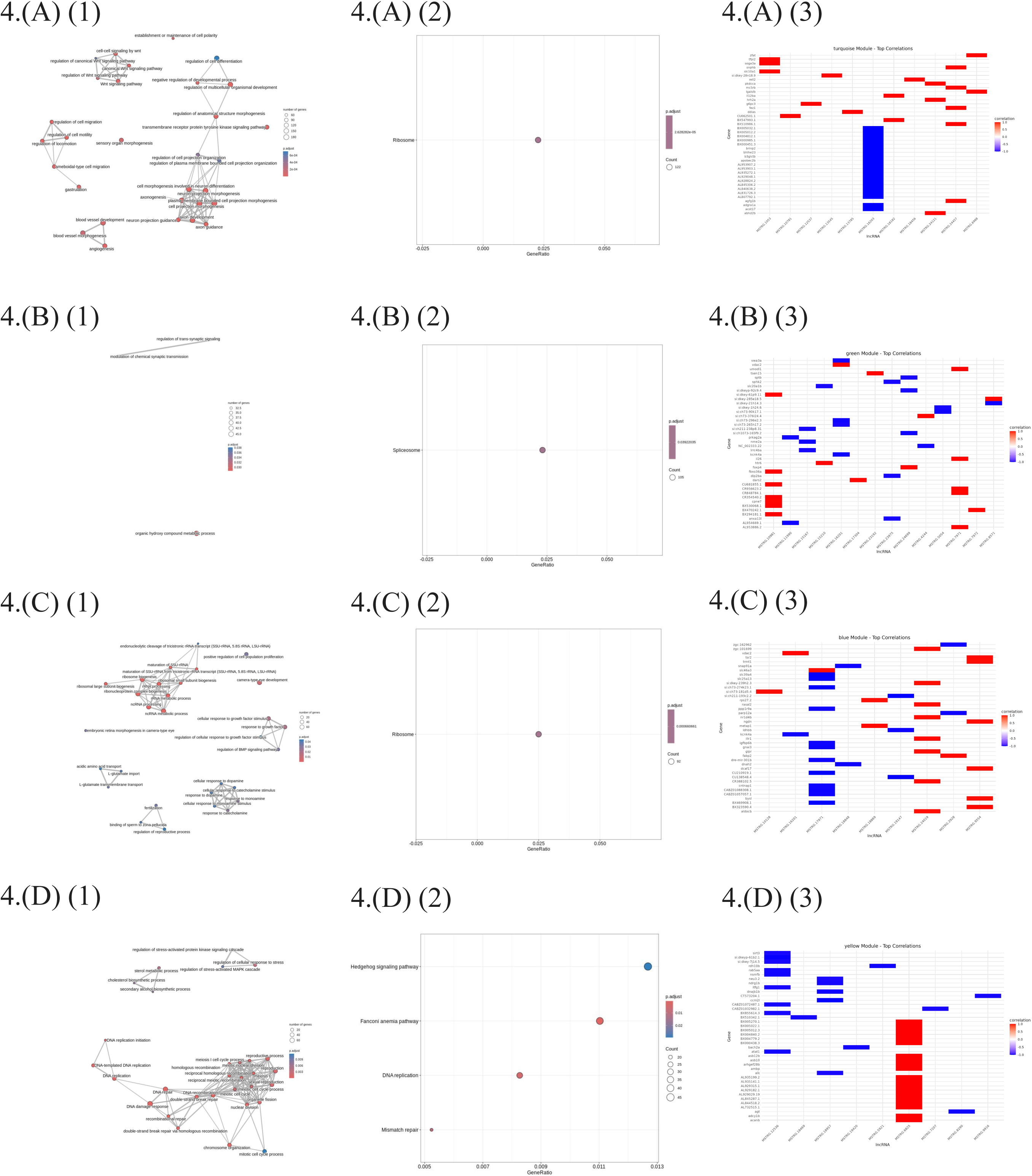
Functional enrichment analysis of gene modules correlated with long non-coding RNAs (lncRNAs). Panels (A) to (D) correspond to the turquoise, green, blue, and yellow modules, respectively. For each module: (1) EMAP plot showing GO enrichment of biological processes for genes with a 90% correlation in expression with lncRNAs, (2) Dot plot of KEGG pathway enrichment analysis for genes highly correlated (r > 0.90) with lncRNAs; (3) The highest and lowest 20 correlation genes with novel lncRNAs. These visualizations highlight the key biological functions and pathways associated with each long non-coding RNA (lncRNA) co-expression module.

These modules exhibited unique expression signatures, each playing a critical role in fin regeneration. Pathway enrichment analysis indicated that lncRNAs in the blue module govern the BMP signaling pathway, ribosome biogenesis, and cellular responses to growth factor stimuli. Turquoise module lncRNAs were enriched in the Wnt signaling pathway and cell projection morphogenesis. The yellow module was linked to DNA damage response, double-strand break repair, and DNA recombination pathways. Finally, lncRNAs within the green module were implicated in the metabolic processes of organic hydroxy compounds.

### Blue Module: BMP Signaling, Ribosome Biogenesis, and Growth Factor Responses

The blue module encompasses critical processes for fin regeneration, including BMP signaling, ribosome biogenesis, and cellular responses to growth factors. Bone Morphogenetic Proteins (BMPs), part of the TGF-β superfamily, play a crucial role in balancing growth and differentiation during zebrafish fin regeneration (R. N. Wang et al., 2014). Disruption of BMP signaling impairs fin growth and differentiation, demonstrating the high importance of regeneration (Smith et al., 2006). MYC oncoprotein drives ribosome biogenesis during regeneration, boosting protein synthesis to support tissue regrowth and differentiation.. These pathways coordinate cellular responses that are integral to tissue repair and regrowth; they control cell proliferation, differentiation, and survival in cells committed to the rebuilding of fins. Although there were no specific citations for fin regeneration studies, TGF-β has been recognized as crucial in tail regeneration contexts, further validating its general relevance in regenerative biology (Ho & Whitman, 2008).

MSTRG.24019.1 exhibits strong positive correlations with three genes involved in regenerative processes. Illr1 (Interleukin-1 Receptor Like 1), correlated at (0.9947124), is necessary for immune responses that support tissue repair (Sebo et al., 2022). aldocb, exhibiting a correlation of (0.9940187), is involved in tissue regeneration (X. Li, 2021). Finally, zgc:101699 (Zebrafish Gene Collection clone 101699) correlates at (0.9921206) and is known to be involved in both regenerative processes and regenerative signaling (Woods et al., 2005). These associations point to a potential role for MSTRG.24019.1 in coordinating immune-mediated and molecular mechanisms vital to effective regeneration.

### Turquoise Module: Wnt Signaling and Cell Projection Morphogenesis

The Wnt signaling pathway plays a pivotal role in zebrafish fin regeneration, acting as a key regulator of cellular processes essential for tissue regrowth and patterning. Wnt/β-catenin signaling defines organizing centers that orchestrate the growth and differentiation of the regenerating fin (Walczyńska et al., 2023). Interestingly, Wnt signaling is not directly active in most proliferative blastemal cells or the epidermis but rather in the nonproliferative distal blastema, suggesting an indirect regulation of cell proliferation and differentiation through secondary signals (Wehner et al., 2014). The pathway’s influence extends to epidermal patterning and osteoblast differentiation, highlighting its role in bone regeneration within the fin (Wehner et al., 2014). Recent studies have shown that both canonical and non-canonical Wnt pathways are involved in various regenerative processes across different species, including zebrafish (Rochard et al., 2016). Cell projection morphogenesis, also regulated by Wnt signaling, is critical for the proper alignment and organization of cells during regeneration, ensuring correct tissue formation and integration (Walczyńska et al., 2023).

MSTRG.1053.1 exhibits significant correlations with three key genes, soga3a, tfpi2, and slc10a1, participating in regenerative processes and tissue maintenance. Soga3a, which shows the strongest correlation (0.9992015), is involved in regenerative processes and contributes to the cellular response to stress (Meng et al., 2017). Tfpi2 (0.9965104) plays a critical part in tissue remodeling, including wound healing and regeneration (Y. Zhang et al., 2013). Similarly, slc10a1 (0.9954736) is linked to cellular homeostasis and supports regenerative responses (Qiu et al., 2017). These associations underline the importance of MSTRG.1053.1 in pathways involved in cellular repair, regeneration, and general tissue health.

### Yellow Module: DNA Damage Response and Repair

The lncRNAs in the yellow module show correlation with genes involved in the DNA damage response (DDR), double-strand break (DSB) repair, and DNA recombination pathways. These pathways play important roles in maintaining genomic stability during the fast cell proliferation and differentiation that take place in fin regeneration. Homologous recombination (HR) accurately repairs DSBs during cell division, preserving genetic integrity, while non-homologous end joining (NHEJ) is faster but error-prone in the absence of templates (Lieber, 2010) (Kieffer & Lowndes, 2022). Recent studies show particle radiation affects DNA end processing and repair, processes crucial for maintaining cell function and enabling regeneration (van de Kamp et al., 2021). While specific studies on fin regeneration were not directly cited, the general importance of these pathways in regeneration is supported by their role in cellular recovery and tissue regeneration across various species (Blanpain et al., 2011).

MSTRG.6825.1 is significantly correlated with three genes of critical importance for regeneration and cellular homeostasis. Specifically, it is significantly negatively correlated (−1) with acanb (actin-related protein), which is key in tissue remodeling, regeneration, and cytoskeleton organization, and adcy1b (adenylate cyclase 1b), an enzyme participating in signal transduction and playing a crucial role in regenerative processes (Schnieder et al., 2020). Further, MSTRG.6825.1 has a correlation with asb10 (Ankyrin repeat and SOCS box containing 10), which is involved in protein degradation pathways, maintaining protein homeostasis, and stress responses during regeneration (Andresen et al., 2014). Together, these correlations point to the possible involvement of MSTRG.6825.1 in the regulation of regenerative pathways and maintaining cellular balance.

### Green Module: Metabolic Processes of Organic Hydroxy Compounds

The green module implicates lncRNAs in the metabolic processes of organic hydroxy compounds, which play significant roles in cellular metabolism and signaling pathways crucial for tissue regeneration (K. C. Wang & Chang, 2011). These compounds are involved in synthesizing and modifying biomolecules essential for cellular growth and repair. The tricarboxylic acid (TCA) cycle, a central metabolic pathway, intersects with the metabolism of organic hydroxy compounds, providing energy and precursors for biosynthetic processes critical during regeneration (Martínez-Reyes & Chandel, 2020). The selective hydroxylation of steroids involves organic hydroxy compounds - a process that may be crucial for fin regeneration, where proper tissue growth and function are achieved by appropriate biochemical modification. Metabolic pathways supply the energy and building blocks essential for cell proliferation and differentiation during regeneration, underscoring their importance in fin repair (Dreher et al., 2020).

Three critical genes involved in regeneration have marked negative correlations with MSTRG.15167.2 . Nme2a (Nucleoside Diphosphate Kinase 2a), with a correlation of (−0.9938864), plays a central role in cell proliferation and regenerative processes (Ferrucci et al., 2024). Lrrc4ba (Leucine Rich Repeat Containing 4Ba), with a correlation of (−0.9902127), is important to neural regeneration; hence, this might suggest MSTRG.15167 plays a role in nervous system repair (Steadman & Sumner, 2018). In addition, si:ch211-238p8.31 (correlation is at −0.9897972) is a zebrafish-specific gene that has been reported to be involved in tissue regeneration (Issaka Salia & Mitchell, 2020). Overall, the negative correlations point towards a possible regulatory function of MSTRG.15167.2 for different regenerative pathways.

### 3.2. Genome-wide discovery and analysis of novel lncRNAs in zebrafish fins, revealing positional memory

The RNA sequencing data, sourced from the Gene Expression Omnibus (GEO) database (accession no. GSE92760), consisted of 14 samples (we failed to download proper SRR5125781_Mid_4 data ) collected from three distinct regions along the proximodistal axis of the zebrafish caudal fin: proximal, middle, and distal. Each region was represented by five biological replicates, with each replicate comprising pooled fin tissue from two male and two female zebrafish. This dataset captures the spatially regulated transcriptomic landscape of the caudal fin, providing insights into the molecular basis of positional memory.

The proximal region is characterized by gene expression patterns associated with enhanced regenerative capacity, while the distal region exhibits signatures linked to positional identity maintenance. The middle region serves as a transitional zone, reflecting a mix of both proximal and distal transcriptomic signatures. Notably, lncRNAs emerge as key regulators in defining positional identity, contributing to spatially coordinated regenerative processes. This classification allows for a comprehensive investigation of the molecular mechanisms underlying positional memory and the spatially distinct regulatory networks that drive zebrafish caudal fin regeneration.

**Transcriptome Assembly and Mapping:**

FastQC showed high-quality sequencing data (Phred >30). After trimming with Trimmomatic, HISAT2 achieved ∼94% mapping to the zebrafish genome (GRCz11). StringTie assembly identified 975,355 transcripts (known and novel), refined to 973,078 after removing unplaced scaffolds.

**Identification of potential LncRNA candidates:**

Using FEELnc, we identified 23,778 putative lncRNAs, of which 10,040 showed non-zero conservation. By selecting transcripts with at least 50% overlap in conserved regions and isolating the top 5% based on conservation metrics, we obtained 3,352 highly conserved lncRNAs, making them strong candidates for functional relevance. PCA of their expression across proximal, middle, and distal fin regions showed that distal samples clustered separately from proximal and middle regions, suggesting a distinct transcriptomic identity and reinforcing the role of lncRNAs in maintaining positional memory during regeneration.

This pattern was further supported by a sample-to-sample distance heatmap, where lower correlations (blue) were observed between the distal region and the other two regions, highlighting its distinct gene expression landscape. These observations suggest that lncRNAs are particularly crucial in defining the positional identity of the distal region, potentially regulating region-specific regenerative processes in zebrafish caudal fin regeneration (Figure 5). Weighted Gene Co-expression Network Analysis (WGCNA) identified 6 modules with unique expression profiles across the fin injury recovery timeline. (Figure 5)

**Figure 5.**
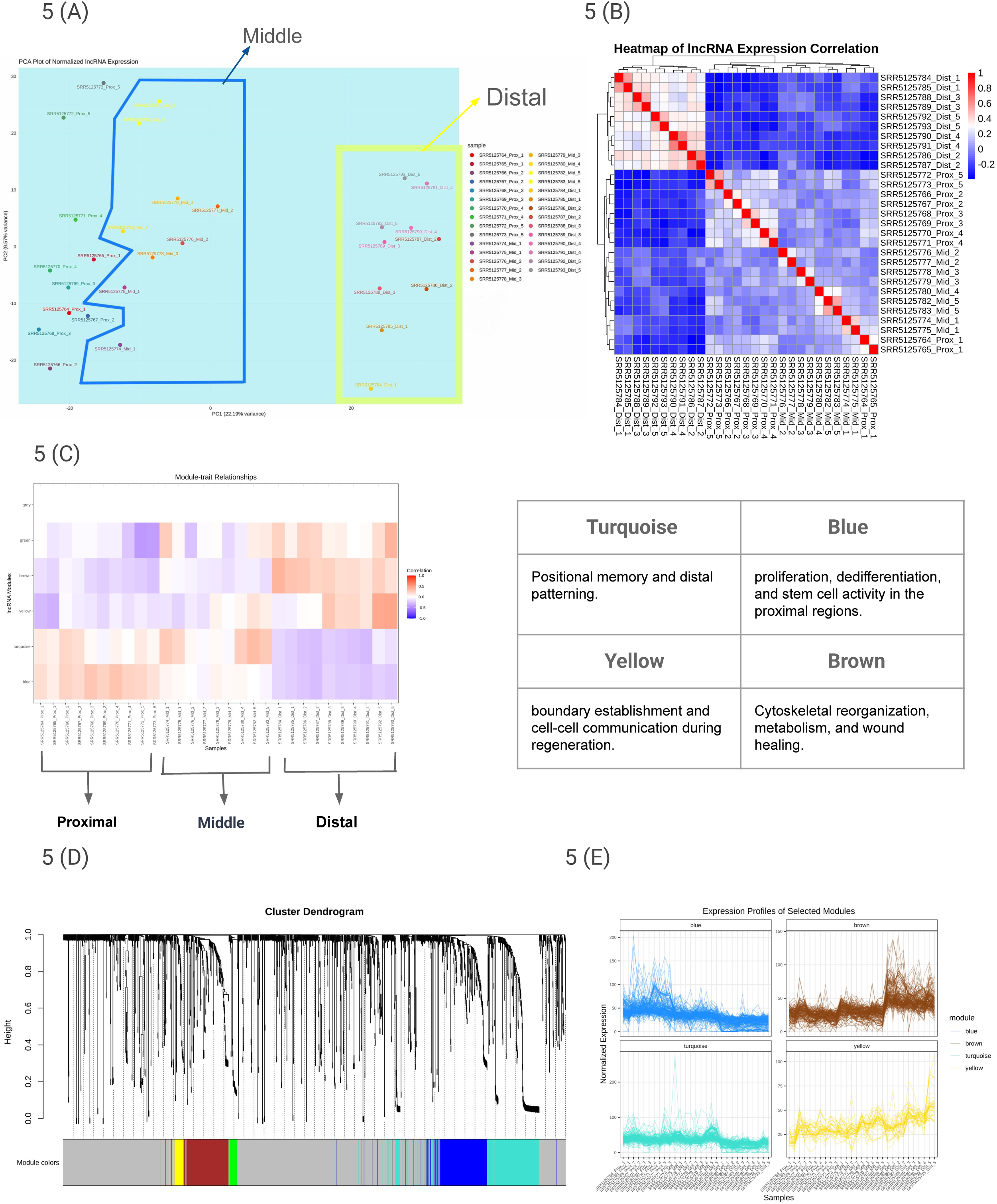
(A) PCA plot of normalized lncRNA expression. (B) Heatmap of lncRNA expression correlation between all samples. (C) Heatmap illustrating the correlation of expression profiles among long non-coding RNA (lncRNA) clusters. Turquoise, Brown, Yellow, and Blue modules show some significant changes. (D) Dendrogram of WGCNA Clusters with 3,352 novel lncRNAs. (E) Expression profiles of the individual long non-coding RNAs in the selected modules co-expressed cluster. The Y-axis indicates normalized gene expression, and the X-axis indicates samples labeled according to the treatment.

Among these modules, 136 novel lncRNA genes were present in the turquoise module, 138 in the blue module, 24 in the yellow module, and 126 in the brown module. Further filtration based on the presence of genes within 2.5 kb of upstream and downstream, with 58 lncRNAs in the turquoise module, 60 in the blue module, 5 in the yellow module, and 57 in the brown module. We further analysed the expression correlation of these lncRNAs with mRNAs and filtered those with r>0.7. The yellow and blue modules failed to correlate with at least 70% correlation in the pathway enrichment process.

**Table 2:**
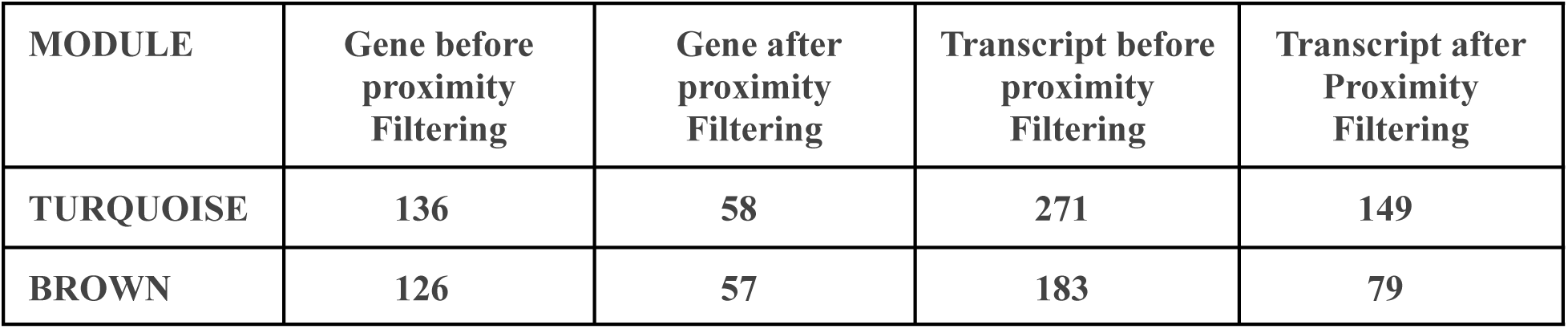
Gene proximity analysis filtering table.

| MODULE | Gene before proximity Filtering | Gene after proximity Filtering | Transcript before proximity Filtering | Transcript after Proximity Filtering |
| --- | --- | --- | --- | --- |
| TURQUOISE | 136 | 58 | 271 | 149 |
| BROWN | 126 | 57 | 183 | 79 |

Gene ontology and pathway enrichment results revealed that lncRNA from brown module showed enrichment for lysosome activity, autophagy, and focal adhesion pathways, suggesting its involvement in cellular remodeling, waste clearance, and spatial regulation during regeneration. In contrast, lncRNAs of the turquoise module were associated with extracellular matrix (ECM) organization, collagen metabolic processes, and bone morphogenesis, highlighting its role in maintaining axial patterning and structural integrity.

These findings indicate that the molecular coordination of positional identity is essential for guiding precise tissue restoration. By regulating distinct biological processes, the brown and turquoise modules collectively shape the regenerative landscape, ensuring the accurate reconstruction of fin architecture and function across different proximodistal segments. ( Figure 6).

**Figure 6.**
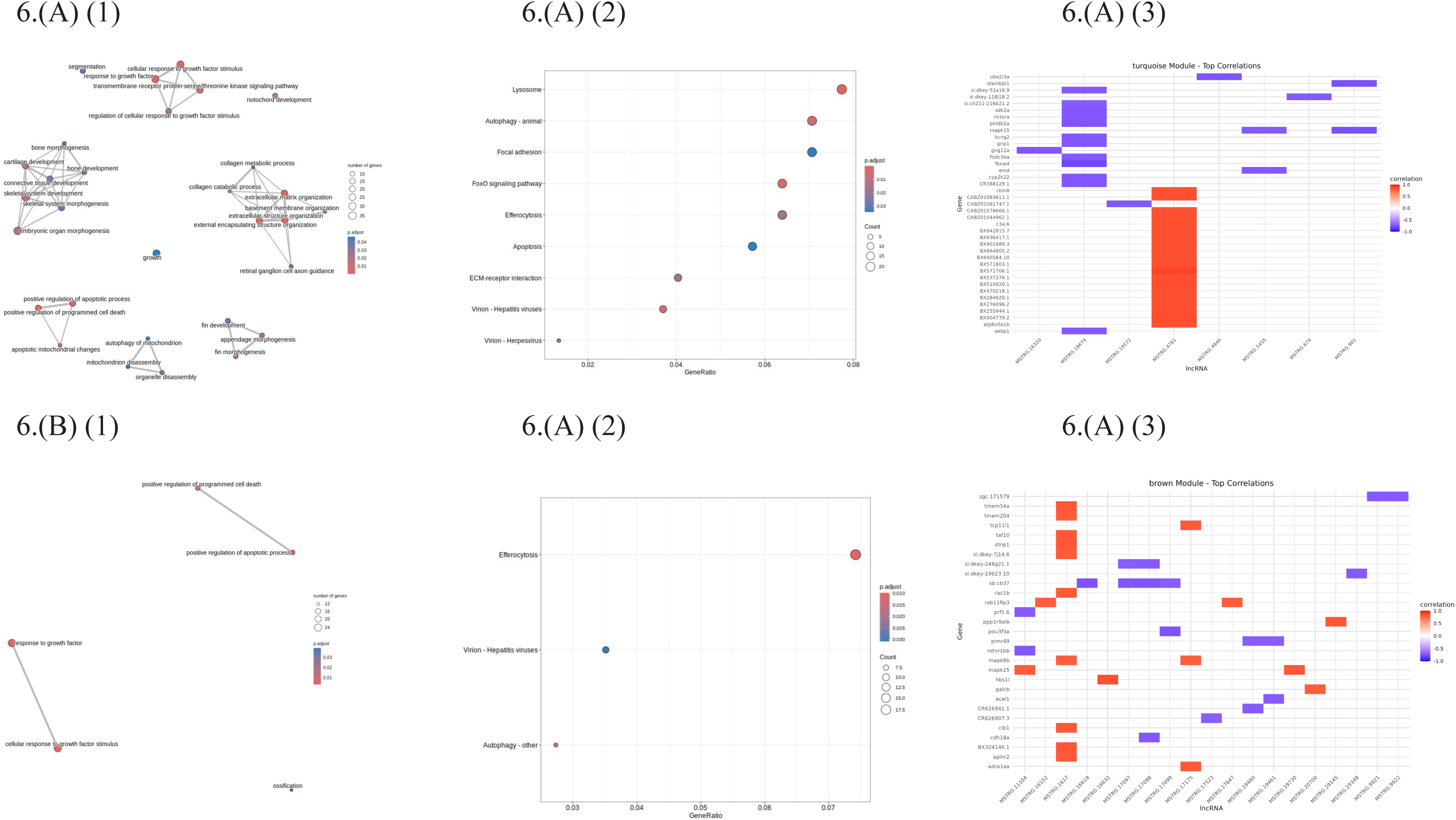
Functional enrichment analysis of gene modules highly correlated with long non-coding RNAs (lncRNAs). Panels (A) and (B) show the turquoise and green modules, respectively. Each panel includes: (1) an EMAP plot depicting Gene Ontology biological processes enriched among genes with expression correlation above 0.70 with lncRNAs; (2) a Dot plot illustrating KEGG pathway enrichment for these highly correlated genes; (3) a list of the top 20 genes with the highest and lowest correlation to novel lncRNAs within each module. These representations offer insights into the biological functions and pathways associated with the identified long non-coding RNA (lncRNA) modules.

### The Brown Module: Regulator of Cellular Remodeling and Positional Memory in Regeneration

The brown module is significantly enriched in pathways related to lysosome activity, autophagy, and focal adhesion, all of which play essential roles in cellular remodeling, waste clearance, and spatial regulation during regeneration (Mizushima et al., 2008) (Levine & Kroemer, 2019). Lysosomes act as metabolic regulators, facilitating the degradation and recycling of cellular waste, which is crucial for maintaining tissue integrity (Saftig & Puertollano, 2021). Autophagy, a tightly regulated process that directs cytoplasmic material to lysosomes for degradation, ensures cellular quality control by eliminating dysfunctional organelles and proteins, thereby supporting regeneration (Klionsky et al., 2021). Additionally, focal adhesion pathways mediate cell-ECM interactions, enabling structural stability and communication necessary for tissue reorganization (Geiger & Yamada, 2011). The interplay between these processes reinforces positional memory, ensuring regenerating cells retain spatial identity by clearing damaged components and restoring their original functions (Mizushima et al., 2008). Collectively, the brown module orchestrates these pathways to facilitate effective tissue repair while maintaining spatial organization essential for precise regeneration.

Analyzing the genes correlated with MSTRG.1617.1 in the brown module, and we identified three key genes showing strong positive correlations that are crucial for regeneration and positional memory: STRIP1 (Striatin Interacting Protein 1) which plays a vital role in cell polarity and cytoskeletal organization essential for positional memory (S. Zhang et al., 2024), TAF10 (TATA-Box Binding Protein Associated Factor 10,) which regulates transcriptional processes during regeneration and helps maintain cell identity (Pahi et al., 2017) and TMEM 54A (Transmembrane Protein 54A) which is involved in membrane organization and cellular patterning during regeneration - together suggesting that this lncRNA coordinates spatial organization, transcriptional regulation, and cellular patterning processes necessary for successful regeneration and positional memory maintenance.

### The Turquoise Module: ECM Organization and Structural Integrity in Regeneration

The turquoise module is strongly associated with extracellular matrix (ECM) organization, collagen metabolic processes, and bone morphogenesis, all of which are essential for maintaining axial patterning and structural integrity during regeneration. ECM organization plays a crucial role in providing biochemical and mechanical support to cells, guiding tissue repair, and structural remodeling (Hynes, 2009). Collagen metabolism is fundamental to ECM stability, influencing cell adhesion, migration, and differentiation, which are critical for regeneration (Kadler et al., 2007). Bone morphogenesis pathways regulate skeletal patterning and repair by modulating osteogenic signals essential for structural reformation (Ortinau et al., 2019). These interconnected processes highlight the turquoise module’s role in orchestrating spatially regulated tissue regeneration, ensuring precise restoration of the zebrafish caudal fin’s architecture and function.

Analyzing the genes correlated with MSTRG.18674.1 in the turquoise module, we identified three key genes showing strong negative correlations that are crucial for regeneration and positional memory: PHLDB2A (Pleckstrin Homology Like Domain Family B Member 2, the correlation ∼ (−0.8)) (H. Li et al., 2024) which is involved in membrane targeting and signaling pathways, GRIP1 (Glutamate Receptor Interacting Protein 1, the correlation ∼ (−0.75)) which plays a role in the synaptic organization and cellular positioning (Dong et al., 1999), and FBXW4 (F-Box And WD Repeat Domain Containing protein, correlation ∼ (−0.75)) which regulates protein degradation during tissue remodeling - together suggesting that this lncRNA may act as a suppressor in coordinating membrane dynamics, cellular positioning, and protein turnover processes necessary for regeneration and positional memory maintenance through negative regulation of these genes (Chen et al., 2024).

**List of Novel lncRNAs :**

The list is given in Supplementary Document 2 for fin regeneration, Supplementary Document 3 for positional memory, we show only two modules of lncRNAs (brown and Turquoise) in positional memory.

### 3.3. Overlapping lncRNAs are crucial to both biological processes

We identified 107 lncRNAs involved in regeneration and 228 lncRNAs associated with positional memory. The regeneration dataset formed four modules: 23 lncRNAs in blue, 46 in turquoise, 18 in yellow, and 20 in green, while positional memory had two modules: 149 lncRNAs in turquoise and 79 in brown. Several overlaps were observed, with 5 lncRNAs from the regeneration blue module overlapping with 4 in the positional memory turquoise module, 8 from the regeneration turquoise module overlapping with 68 in positional memory turquoise, 6 from regeneration turquoise overlapping with 8 in positional memory brown, and 3 from regeneration yellow overlapping with 3 in positional memory brown. The list of overlapping lncRNAs is provided in Supplementary Document 4. After analyzing their positions and identifying the most common regions, we identified 13 key regions where either the full or partial sequence of an lncRNA plays a significant role in both regeneration and positional memory. These findings offer valuable insights into their potential regulatory functions and their contribution to tissue regeneration and spatial identity.

**Table 3.** Main important overlapping region among all lncRNAs, which is important in both biological processes.

| CHR | Start Position | Stop Position | STAND | NAME |
| --- | --- | --- | --- | --- |
| 1 | 59386284 | 59389365 | + | ZEB_RE_POS_FIN_1 |
| 1 | 40651504 | 40659496 | - | ZEB_RE_POS_FIN_2 |
| 1 | 52379492 | 52380379 | - | ZEB_RE_POS_FIN_3 |
| 2 | 18334246 | 18335688 | + | ZEB_RE_POS_FIN_4 |
| 3 | 7463694 | 7479910 | + | ZEB_RE_POS_FIN_5 |
| 4 | 47597472 | 47599262 | + | ZEB_RE_POS_FIN_6 |
| 4 | 62329752 | 62330888 | + | ZEB_RE_POS_FIN_7 |
| 5 | 11497931 | 11500894 | + | ZEB_RE_POS_FIN_8 |
| 7 | 32321510 | 32321813 | - | ZEB_RE_POS_FIN_9 |
| 16 | 54377947 | 54378474 | + | ZEB_RE_POS_FIN_10 |
| 18 | 23459613 | 23461429 | + | ZEB_RE_POS_FIN_11 |
| 19 | 19458863 | 19461118 | - | ZEB_RE_POS_FIN_12 |
| 23 | 5048681 | 5052895 | - | ZEB_RE_POS_FIN_13 |

The spatiotemporal expression dynamics of the 13 selected lncRNAs are illustrated in Figure 7 . Figure 7(C) displays the expression variation of each lncRNA across different regeneration time points, highlighting temporal changes in transcriptional activity during the regenerative process. Figure 7 (D) shows the positional expression differences of the same lncRNAs across the proximal, median, and distal regions of the regenerating fin, emphasizing their region-specific regulation. A more detailed, integrative analysis is presented in Supplementary Document 5, where BAM files from various time points were merged with those from the positional memory dataset and normalized. This combined view demonstrates how each lncRNA’s expression is influenced over time in the context of its spatial identity, providing deeper insights into the coordinated regulation of lncRNAs during regeneration.

**Figure 7.**
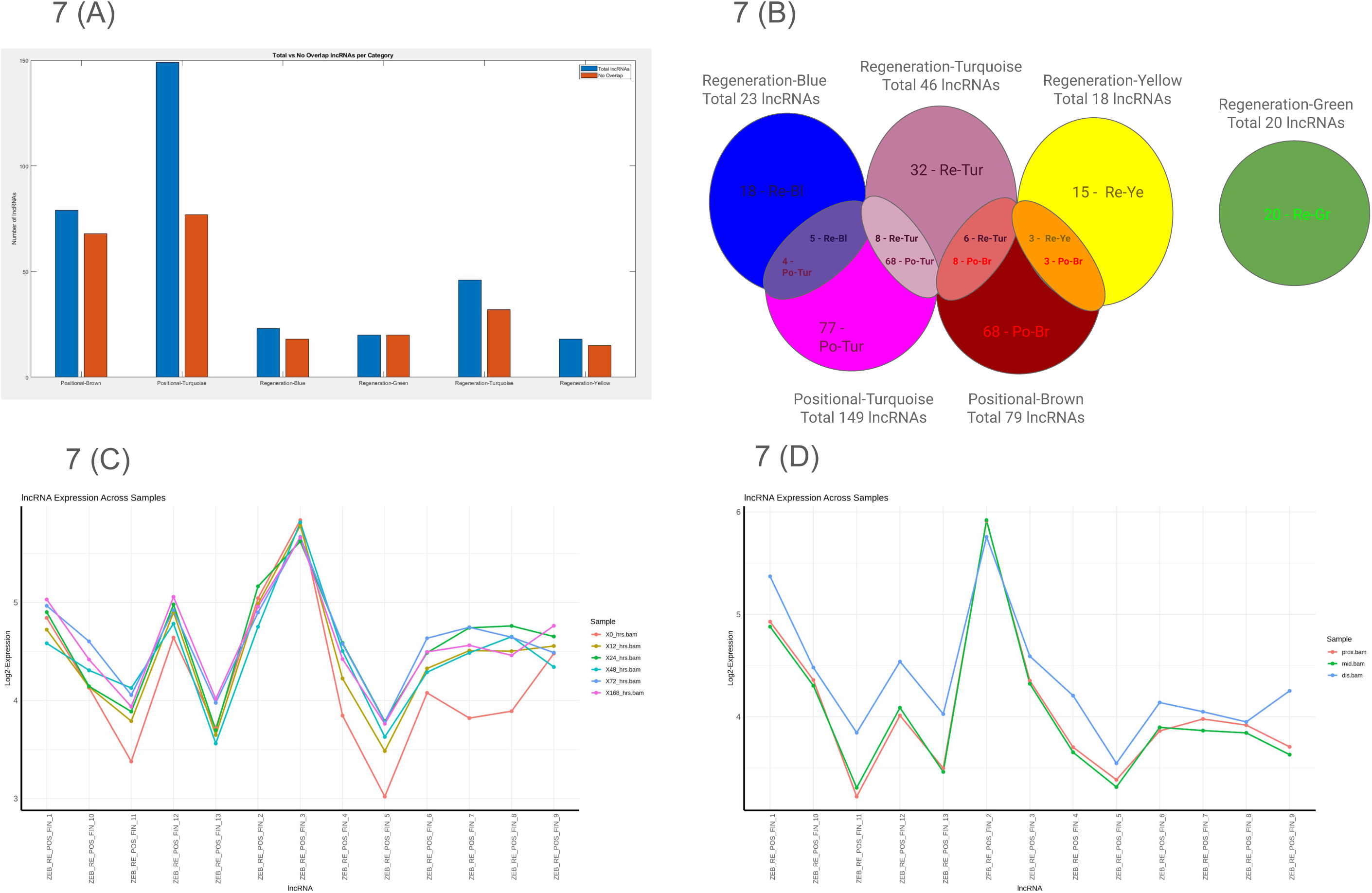
(A) number of total lncRNAs with non-overlapping lncRNAs, (B) how positional memory module lncRNAs overlapped with fin regeneration lncRNAs. For example, six Re-Tur are overlapped with eight Po-Br lncRNAs. (C), Expression varies in different positional points of each 13 lncRNAs, (D) Expression varies in different time points of each 13 lncRNAs.

### 3.4. WGCNA and pathway analysis with overlapping regions

After merging StringTie CSV files from both the regeneration and positional memory RNA-seq datasets, we created a count matrix, which was then normalized based on groups. The pipeline is provided in Figure 8. WGCNA analysis was performed on this normalized matrix, generating a dendrogram with 89 color modules in Supplementary Document 6. We also performed PCA plotting and generated a heatmap to analyze sample-to-sample variations. Figure 9 presents a comprehensive expression analysis, including: A) a PCA plot of normalized RNA expression, B) a heatmap of sample-wise expression correlation, and C) a cluster-level heatmap showing expression correlations among regions containing the 13 overlapping lncRNAs.

**Figure 8.**
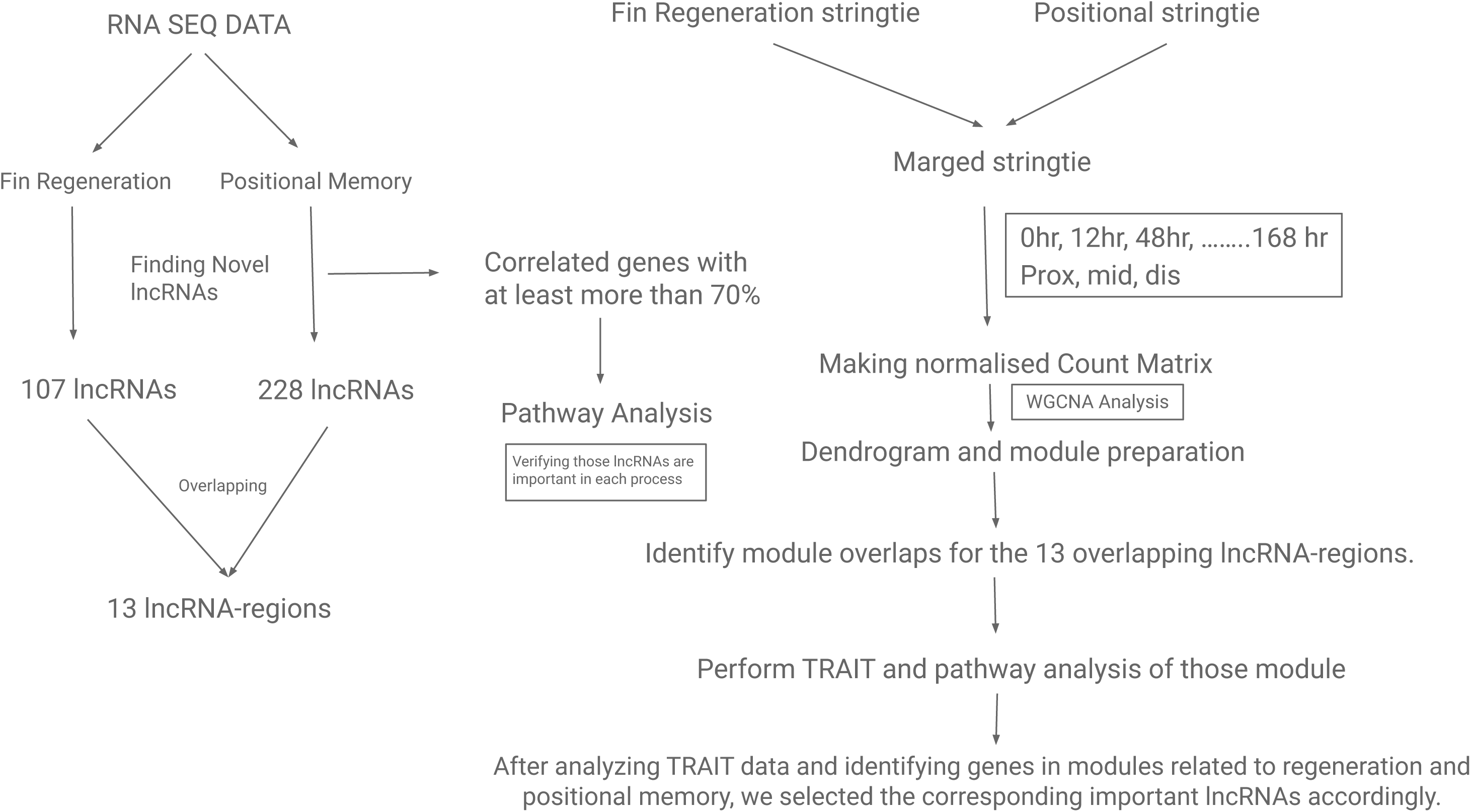
Schematic of the lncRNA verification pipeline

**Figure 9.**
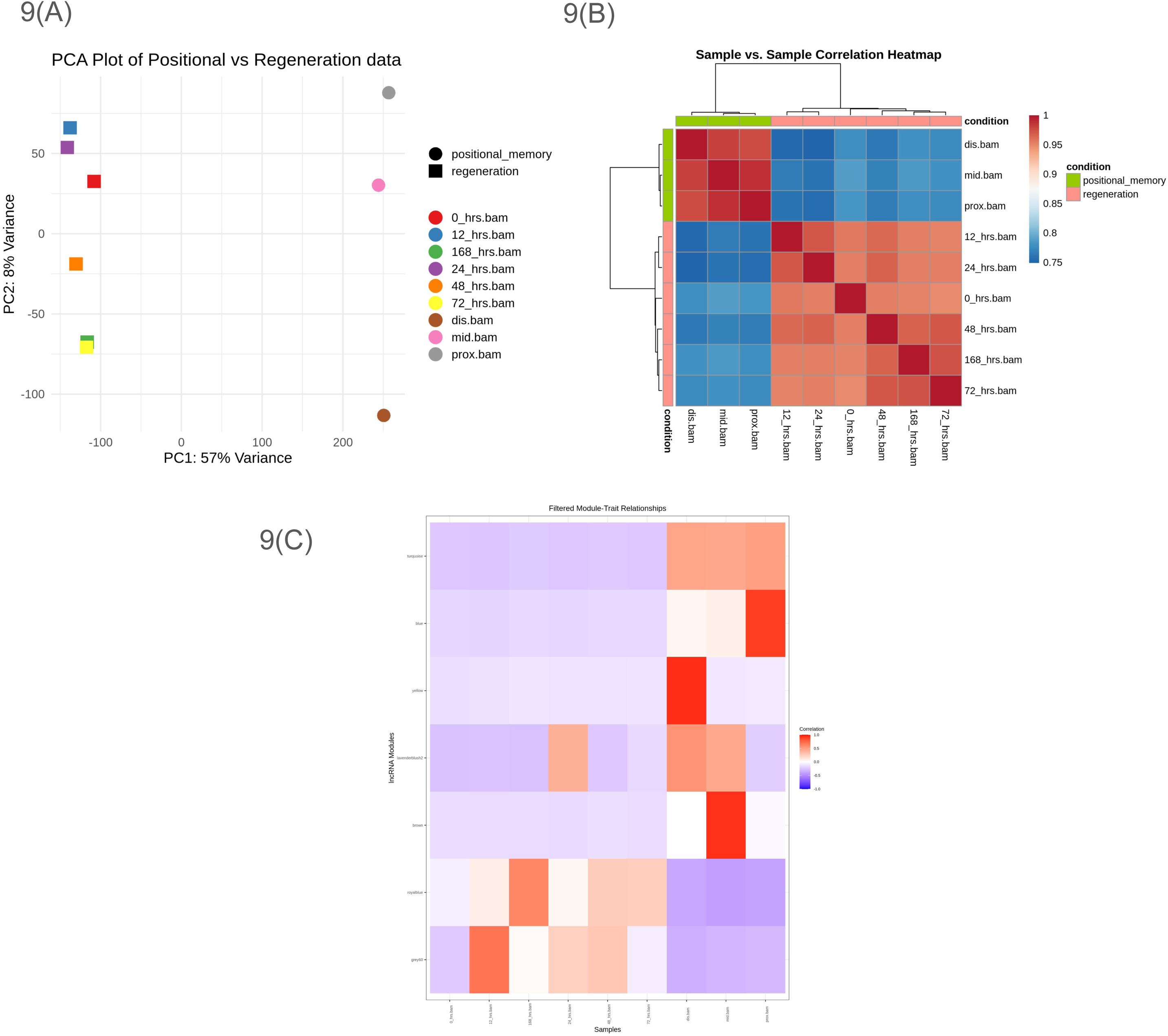
A) PCA plot of normalized RNA expression. B) Heatmap of RNA expression correlation between all samples. C) Heatmap showing the correlation of expression profiles among RNA clusters where all my overlapping 13 lncRNA-regions are present.

Next, we identified lncRNAs that fully overlap with the 13 previously detected overlapping lncRNA regions. Since merging the StringTie files modified the lncRNA names, we reassigned these regions to their corresponding modules. Our analysis confirmed that these regions are present in the turquoise, blue, yellow, lavenderblush2, brown, royal blue, and grey60 modules.

Finally, we performed TRAIT analysis to assess the significance of these modules. The results revealed that the brown, lavender blush 2, royal blue, and grey 60 modules exhibited a strong and noteworthy trend. We then conducted pathway enrichment analysis, and the enrichment maps (EMap) are provided in Supplementary Document 7. The study identifies many important pathways. Recent research has highlighted the intricate interplay between various molecular processes in tissue regeneration. Chromatin remodeling and epigenetic modifications play crucial roles in regulating DNA repair mechanisms, which are essential for maintaining genome stability during regenerative processes (Leiba et al., 2023) (Guo et al., 2022). The dynamic regulation of chromatin accessibility, influenced by nucleosome positioning and epigenetic modifications, is key to facilitating efficient DNA recombination and repair (Klemm et al., 2019). Additionally, specific lipid signaling pathways and metabolic shifts have been shown to modulate regeneration in various tissues, including the liver and neural tissues (Dakal et al., 2025). The epidermal growth factor receptor (EGFR) and ErbB signaling pathways are also critical in tissue regeneration, regulating cellular processes such as proliferation, differentiation, and survival (Dakal et al., 2025). These pathways are particularly important in maintaining tissue integrity and promoting repair processes in organs like the intestine and liver (Mayorca-Guiliani et al., 2025). Understanding the complex interactions between these pathways and processes is crucial for developing targeted therapeutic strategies to enhance tissue repair and regeneration in clinical settings.

In the lavenderblush2 module, the most crucial gene is DNase1. DNase1 plays a role in apoptosis and tissue remodeling, facilitating the removal of damaged cells and shaping new tissue grow (Keyel, 2017) (Martínez Valle et al., 2008). TTLL9 is involved in microtubule modifications, particularly tubulin polyglutamylation, which is essential for cellular reorganization during regenerative processes (Jentzsch et al., 2024) (Genova et al., 2023). FAM50A has been implicated in transcriptional regulation, potentially influencing the expression of genes necessary for regeneration, including its role in the spliceosome complex and modulation of cell proliferation and apoptosis. PLEKHD1 is thought to play a role in cellular signaling pathways that are involved in tissue repair and regeneration, potentially interacting with or influencing pathways similar to the well-studied Wnt/β-catenin and Notch signaling cascades (Liu et al., 2022) (Nair et al., 2019). The specific roles of CR855375.1, BX324179.1, CR388363.1, and CR847944.1 (Howe, 2001) in regeneration are currently undocumented. Their presence in the lavenderblush2 module, a co-expressed gene network potentially active during regeneration, suggests possible involvement in regenerative processes (Seifert et al., 2023). These modules of lncRNAs may be involved in crucial biological processes such as cell proliferation, apoptosis, angiogenesis, and extracellular matrix remodeling, which are important in regeneration (Fernández-Guarino et al., 2023). This presents an opportunity for targeted studies to elucidate their functions in tissue repair and regeneration (Xia et al., 2018).

In the royal_blue module, the regenerative process involves a complex interplay of various genes, each contributing to specific aspects of tissue repair and renewal. Key players include id1, crucial for cell cycle regulation and stem cell maintenance (Echeverri & Zayas, 2018), and hmgb1b, a DAMP protein facilitating wound healing (Jin et al., 2023). Manf supports cell survival and tissue repair, while mafba contributes to development and regeneration (Jin et al., 2023). Annexins (anxa1c and anxa5b) regulate inflammation and apoptosis, essential for wound healing (Han et al., 2020) . Other significant genes include tk2 and exo5 for DNA repair, pde6gb for signaling, rnf169 for DNA damage repair, laptm4b for cell proliferation, cyp46a1.3 for lipid metabolism, irx6a for tissue development, stat2 for immune signaling, and cul2 for protein degradation and cell cycle regulation (Kaina, 2020). This diverse array of genes highlights the multifaceted nature of regeneration, encompassing processes from cellular repair to tissue-wide renewal. CR381540.3 and BX322787.1 are potentially crucial for zebrafish regeneration, influencing cell proliferation and inflammatory responses (Taylor et al., 2024). These modules of lncRNAs may regulate gene networks essential for wound healing and blastema formation, key processes in regeneration (Mattick et al., 2023). Further research is needed to elucidate their specific roles in regenerative signaling and cellular reprogramming, which could have implications for understanding tissue repair mechanisms (Katsuyama & Paro, 2011).

For the grey60 module, the regenerative process involves a complex interplay of genes regulating various cellular mechanisms. Adam10a, casp6b.1, and srgn are crucial for tissue remodeling and repair, with roles in processes such as apoptosis, inflammation, and extracellular matrix remodeling (Naba, 2024). Arid2 influences gene expression essential for regeneration, particularly in stem cell differentiation and osteoblast commitment (Xu et al., 2012). Zbed4, rps9, and rab5b likely contribute to cell proliferation, protein synthesis, and cellular reprogramming, respectively, based on their known functions in related cellular processes (Saghizadeh et al., 2011) . Ube2l3b, as part of the ubiquitin-proteasome system, regulates protein degradation in regenerative signaling pathways, while tmem161a facilitates cell communication during regeneration through its role in signal transduction (Ji et al., 2019). This module contains several lncRNAs previously known for their importance in regeneration, including H19, Linc-MD1, LncMyoD, Malat1, and SRA, which are primarily involved in muscle differentiation, tissue repair, and stem cell regulation (Statello et al., 2021). The specific roles of BX548015.2, CU019562.3, CT027980.1, CT573366.1, BX088688.2, BX936391.1, CU681836.1, AL954138.2, and CR792438.1 (Howe, 2001) in regeneration are currently undocumented. Their presence in the grey_60 module, a co-expressed gene network potentially active during regeneration, suggests possible involvement in regenerative processes (Montenegro, 2022). Additionally, this module includes lncRNAs that could play a significant role in regeneration, which require further investigation to determine their precise functions in wound healing, blastema formation, and cellular reprogramming (Mattick et al., 2023).

The brown module contains several genes associated with regeneration, with some key lncRNAs playing crucial roles. Hypoxia-inducible factor 1-alpha (HIF-1α) regulates angiogenesis and osteogenesis by elevating VEGF levels in osteoblasts. Transforming growth factor-beta 1 (TGF-β1) signaling, activated under hypoxic conditions via HIF-1α, promotes collagen deposition, crucial for tissue repair. Additionally, HIF-1α is essential for bone regeneration, influencing various pathways in bone tissue engineering (You et al., 2023) (You et al., 2023). Additionally, CR381540.2 has been linked to cellular differentiation processes, which are vital for tissue regeneration, while CR788322.2 is involved in regulating gene expression during stress responses, potentially influencing regenerative outcomes. These modules of lncRNAs present significant opportunities for understanding molecular mechanisms in regeneration and warrant further investigation.

After WGCNA analysis and pathway enrichment, we identified four key lncRNA regions among the 13 overlapping lncRNA regions that play a major role: ZEB_RE_POS_FIN_2, ZEB_RE_POS_FIN_4, ZEB_RE_POS_FIN_5, and ZEB_RE_POS_FIN_11. These regions are distributed across multiple modules, with ZEB_RE_POS_FIN_11 present in grey60 and turquoise, ZEB_RE_POS_FIN_2 in brown, blue, and turquoise, ZEB_RE_POS_FIN_4 in lavenderblush2, and ZEB_RE_POS_FIN_5 in royal blue and turquoise. The list of all regions and their corresponding modules is provided in Supplementary Document 8.

Figure 7 (C) illustrates the expression variation of each lncRNA across different regeneration time points, highlighting temporal changes in transcriptional activity during the regenerative process. Figure 7 (D) shows the positional expression differences of the same lncRNAs across the proximal, median, and distal regions of the regenerating fin, emphasizing their region-specific regulation.

### 3.5 Validation of Selected lncRNAs

To validate our in silico findings, we selected the eight most effective lncRNA regions ZEB_RE_POS_FIN_1, ZEB_RE_POS_FIN_2, ZEB_RE_POS_FIN_4, ZEB_RE_POS_FIN_5, ZEB_RE_POS_FIN_9, ZEB_RE_POS_FIN_11, ZEB_RE_POS_FIN_12, and ZEB_RE_POS_FIN_13 that displayed strong and stage-specific expression in the RNA-seq dataset. The qRT-PCR results confirmed that all eight regions exhibited clear and consistent expression at the specific regeneration time points where they were predicted to be active. This alignment between experimental and computational data strengthens the biological relevance of these long non-coding RNA (lncRNA) regions. The validated expression patterns are presented in Figure 10, further supporting their potential role in fin regeneration. The results confirmed the stage-specific and dynamic regulation of these lncRNAs (Figures 7 and 10).

**Figure 10.**
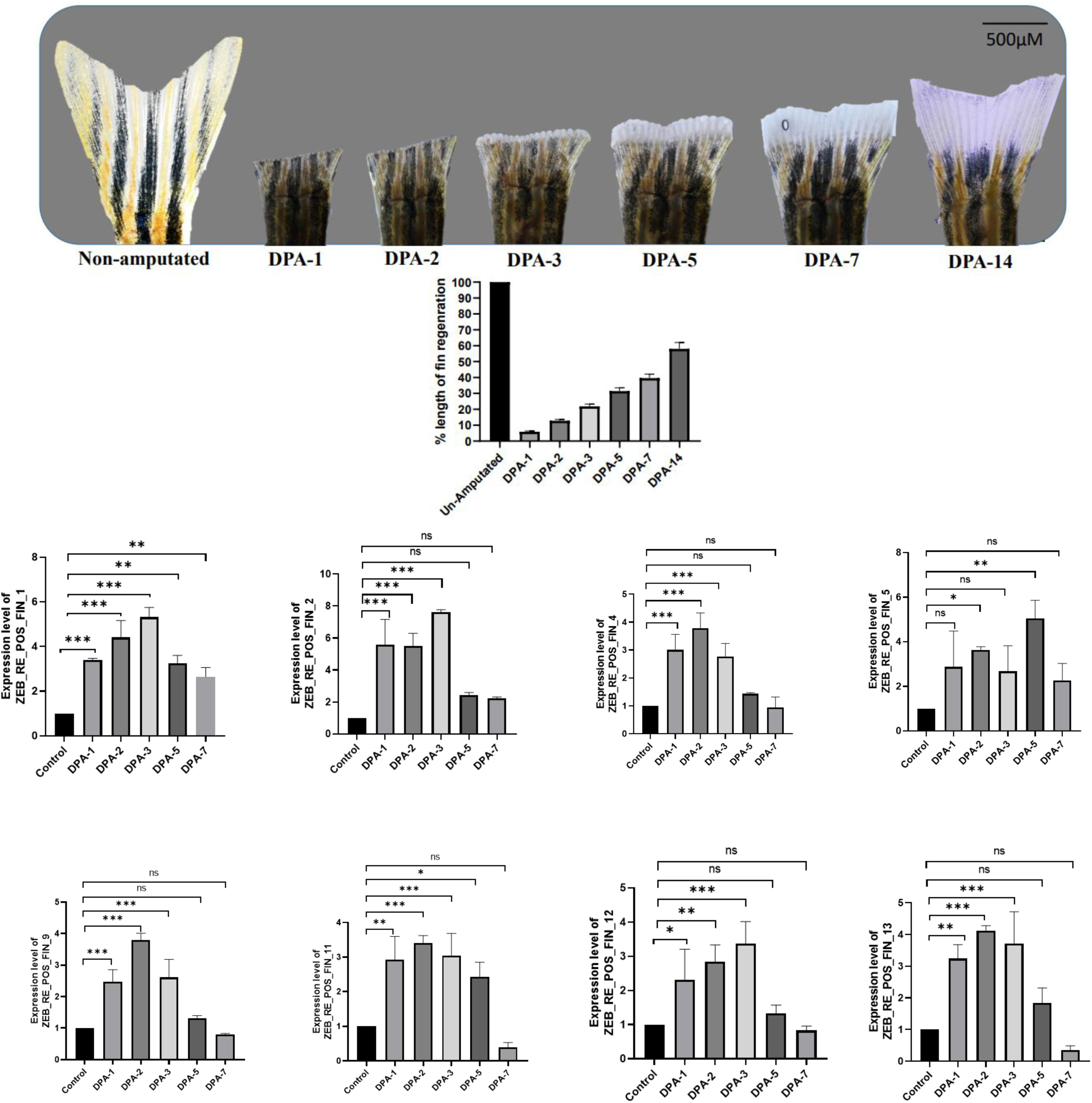
Expression of lncRNA-region in Zebrafish Caudal Fin Regeneration in Proximal Position.

**Figure 11.**
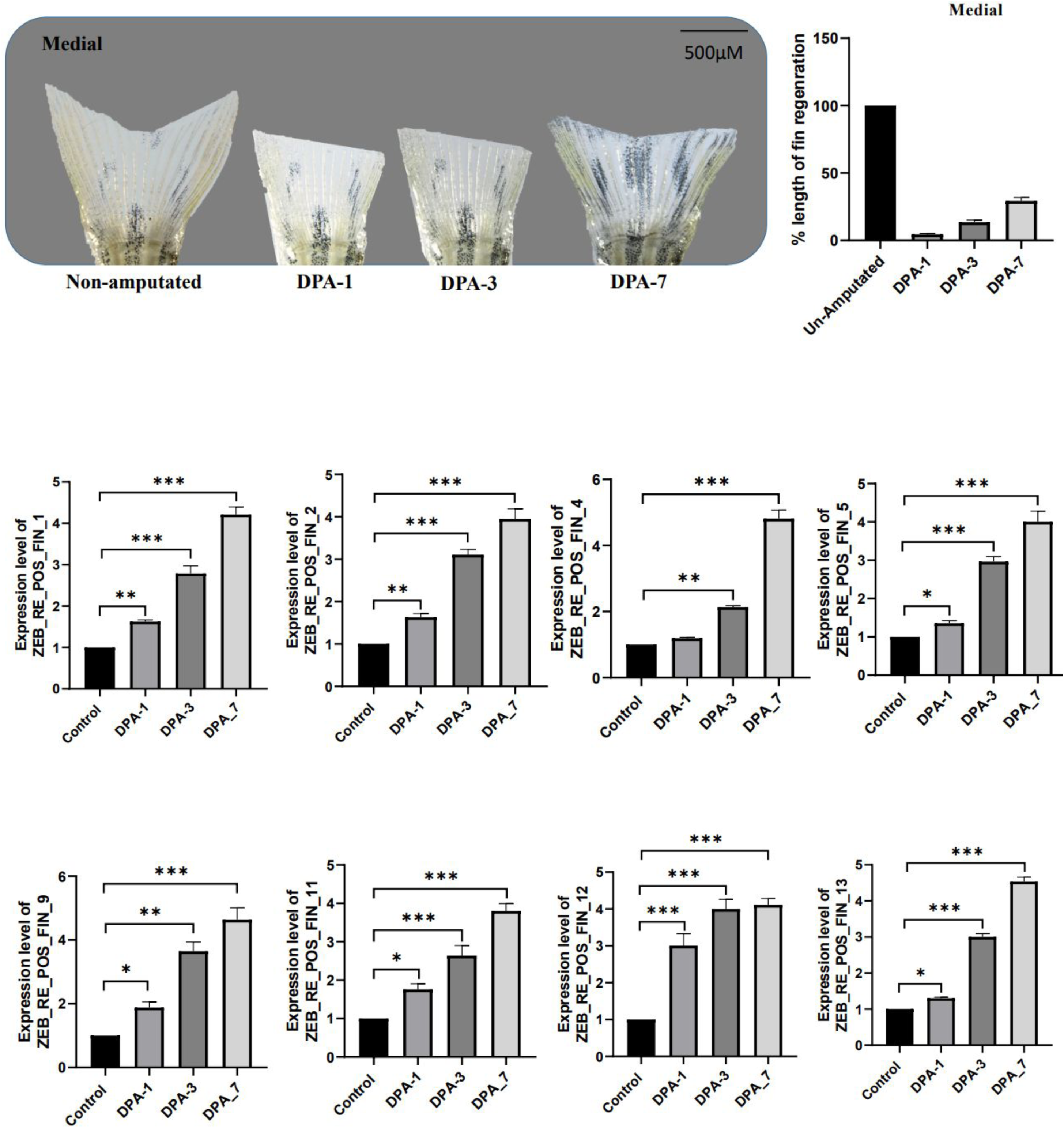
Expression of lncRNA-region in Zebrafish Caudal Fin Regeneration in Medial Position.

**Figure 12.**
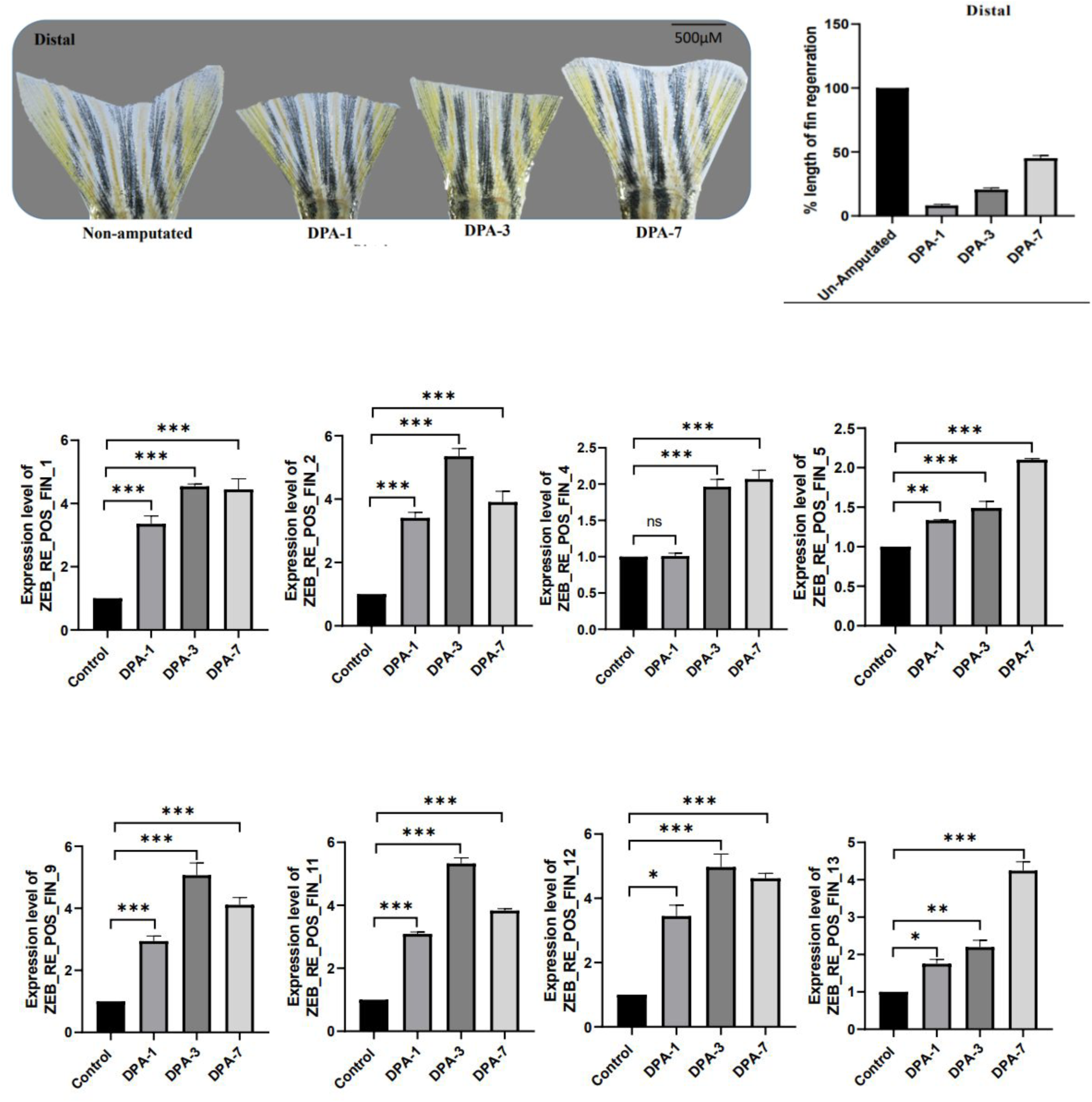
Expression of lncRNA-region in Zebrafish Caudal Fin Regeneration in Distal Position.

**Early induction (1–2 dpa):** *ZEB_RE_POS_FIN_1, _2, _4, _9, and _11* were strongly upregulated as early as 1 dpa, peaking between 2-3 dpa. For example, *ZEB_RE_POS_FIN_1* showed ∼5-fold induction at 2 dpa, while *ZEB_RE_POS_FIN_2* and *ZEB_RE_POS_FIN_9* displayed ∼7-8-fold upregulation compared to the control. These profiles suggest a role in wound healing, blastema formation, and early proliferative events.

**Intermediate/later activity:** *ZEB_RE_POS_FIN_5* and *ZEB_RE_POS_FIN_12* showed modest but significant induction at 2-3 dpa (∼2–3 fold), indicating possible roles in the transition from proliferation to differentiation. *ZEB_RE_POS_FIN_13* peaked at 3 dpa (∼5-fold), declining thereafter, consistent with functions in mid-regeneration events.

**Decline at remodeling stage (7 dpa):** Most lncRNAs (ex, *ZEB_RE_POS_FIN_1, _2, _4, _9, _11, _13*) showed reduced expression by 7 dpa, suggesting their functions are temporally restricted to early and intermediate regeneration phases.

Overall, the temporal expression profiles demonstrated a strong concordance between computational predictions and experimental validation (Figure 7A). Their expression dynamics highlight the importance of further investigation into these lncRNA regions, as they were identified through PhastCons conservation analysis and also demonstrate positional memory characteristics. Together, these findings suggest that these lncRNAs may play critical roles in regenerative processes and could potentially serve as conserved regulators of regeneration in other species.

To support the positional relevance of our identified lncRNAs across multiple species, we performed qPCR on days 1, 3, and 7 in both the proximal, median, and distal regions of the regenerating tissue (Figures 10, 11, and 12). The results revealed distinct expression patterns at each position during the time point, which indicates each early induction, Intermediate activity, and remodeling stage, indicating that these lncRNAs are not only position and time specific, but may also play important roles in evolutionary conserved regenerative processes.

ZEB_RE_POS_FIN_1 exhibited a marked downregulation by day 7 in the proximal fin region, whereas in the median and distal regions, its expression peaked around day 3 of regeneration, suggesting a region-specific temporal expression pattern. ZEB_RE_POS_FIN_2 displayed an upward trend in expression in the median fin region, while in the proximal and distal regions, its expression declined significantly after day 3. Notably, expression in the distal region remained consistently low throughout the regeneration timeline. ZEB_RE_POS_FIN_4 showed markedly low expression in the proximal region by day 7, while the median region exhibited a consistent upward trend throughout regeneration. Although the distal region also followed an increasing trend, its expression remained relatively low from day 3 onward. ZEB_RE_POS_FIN_5 exhibited a downward trend in the proximal region, while both the median and distal regions showed an upward trend in expression. However, the increase in the median region was comparatively less pronounced. ZEB_RE_POS_FIN_9 shows a peak at Day 3 in the proximal and distal regions, followed by a decline by Day 7, while in the median region, expression steadily increases from Day 1 to Day 7, reaching its highest level at DPA-7. ZEB_RE_POS_FIN_11 exhibited a sharp upregulation by Day 3 in both the median and distal regions, followed by a sustained high or slightly decreased expression by Day 7. In contrast, the proximal region showed an early increase at Day 1 and Day 3, but a sharp drop in expression by Day 7, suggesting a transient activation pattern unique to this region. ZEB_RE_POS_FIN_12 showed a transient expression peak at Day 3 in the proximal region, followed by a clear downregulation by Day 7. In contrast, both the median and distal regions demonstrated a strong and sustained upregulation from Day 1 through Day 7, with no significant drop in expression. ZEB_RE_POS_FIN_13 also exhibited a distinct expression pattern in the proximal region compared to the median and distal regions, indicating region-specific regulation during regeneration.

In summary, the qPCR data not only validate the 3-day expression patterns shown in Figure 7(D), but also reveal clear temporal and positional dynamics of lncRNA expression across the fin during regeneration. Distinct trends were observed across the proximal, median, and distal regions. In the proximal region, lncRNAs displayed a transient expression profile, peaking around Day 2 or 3 and then declining sharply by Day 7, indicating early but short-lived involvement. The median region exhibited a gradual and sustained increase from Day 1 to Day 7, suggesting continuous regulatory activity throughout regeneration. In contrast, the distal region showed robust and consistently high expression levels from early to late stages, reflecting persistent activation. Together, these findings emphasize the spatially and temporally distinct roles of lncRNAs in coordinating the regenerative response across different fin regions.

## 4. Discussion

Our study identifies a set of lncRNA regions that are spatiotemporally regulated during zebrafish caudal fin regeneration, integrating both regeneration time-course and positional memory datasets. By overlapping these datasets, we identified 13 lncRNA genomic regions that are consistently present during key regenerative stages, span the proximodistal axis over time, and are also conserved across multiple models. Experimental validation via qRT-PCR confirmed their stage-specific expression, particularly during blastema formation and regenerative outgrowth, aligning with our computational predictions. These results highlight that lncRNAs serve as dynamic molecular regulators, linking positional identity with regenerative progression.

To identify these novel long non-coding RNAs (lncRNAs), we developed a comprehensive bioinformatics pipeline. RNA-seq reads from both datasets were quality-checked and aligned using HISAT2, and transcripts were assembled with StringTie to create sample-specific transcriptomes. FEELnc was used to filter and classify lncRNAs based on coding potential and genomic context, while WGCNA co-expression analysis linked lncRNAs to relevant gene modules associated with regeneration and positional memory. PhastCons conservation scoring was then applied to prioritize evolutionarily constrained regions. We also recheck the effectiveness of lncrnas-region via doing a full WGCNA analysis with all other RNAs, and final candidates were validated against RNA-seq expression profiles and qRT-PCR results, ensuring that only robust, biologically relevant lncRNAs were selected.

The temporal expression profiles of the validated lncRNAs further reinforce their potential role in coordinating positional memory with regenerative progression. For instance, lncRNAs peaking at early stages (1-2 dpa) overlapped with transcripts enriched in proximal and middle fin compartments, consistent with their association with blastema initiation and proliferative cues. Conversely, those induced at intermediate stages (3-5 dpa) aligned with positional signatures from distal compartments, suggesting involvement in differentiation and patterning processes.

This convergence across datasets underscores that lncRNAs are not only time-specific regulators of regeneration but also spatially tuned, providing a molecular link between positional identity and regenerative outcome.

Importantly, the conservation analysis suggests that several of these lncRNA regions are evolutionarily conserved, raising the exciting possibility of identifying universal lncRNAs that may govern regeneration across species. Exploring these regions in mammals or other vertebrates could uncover latent regenerative mechanisms, potentially enabling the design of lncRNA-based therapeutic strategies. From a future perspective, functional characterization of these candidates through knockdown, overexpression, or CRISPR-based perturbation approaches will be critical to establish their roles in regeneration. Such studies could bridge descriptive profiling with mechanistic insights and may ultimately guide translational efforts in regenerative medicine.

Our findings thus provide a foundation for cross-species studies, moving closer to defining core non-coding RNA regulators that could one day enhance tissue repair and regeneration in species with limited regenerative capacity.

## 5. Data availability

The RNA sequencing data used for the positional memory analysis were obtained from the Gene Expression Omnibus (GEO) under the accession number GSE92760. The RNA-seq data used for the regeneration time-course analysis were retrieved from the NCBI Sequence Read Archive (SRA) under the BioProject accession number PRJNA248169. All processed and intermediate files generated during this study are available from the corresponding author upon reasonable request.

## Acknowledgement

I would like to thank my supervisor and collaborators for their invaluable guidance and support throughout this study. I am also grateful to my lab colleagues for their assistance and constructive feedback, and to the funding agencies that made this research possible.

## Author Contributions

Soumyadeep Paul: Conceptualized and designed the study; curated and analyzed publicly available zebrafish regeneration datasets with a specific focus on positional memory; developed and automated the lncRNA discovery pipeline. Identified novel lncRNAs associated with both regeneration and positional memory, and established a comparative framework to relate positional memory signatures with time-resolved regeneration data to capture dynamic regulatory changes. Performed integrative analyses to identify lncRNAs with shared and distinct roles across spatial and temporal contexts; conducted position- and time-specific expression profiling; and identified key lncRNAs with consistent differential expression across datasets. Validated candidate lncRNAs through correlation with regeneration-associated genes and assessed their potential evolutionary conservation. Designed the qPCR validation strategy, including primer design and fin dissection protocol. Interpreted findings in light of existing literature and prepared the complete manuscript with integrated data, figures, and validation results.

A Hariharan: Maintained and handled zebrafish lines; performed fin amputations following the experimental protocol; extracted RNA, synthesized cDNA, conducted RT-PCR experiments, and provided the resulting expression graphs for analysis.

Dasari Abhilash and Surbhi Kohli: Provided overall guidance and critical input throughout the course of the study.

Dr. Shilpi Minocha: Conceived and supervised the overall project; supported funding and execution of wet-lab procedures; assisted in analysis and interpretation of experimental results, and provided continuous guidance throughout all stages of the study, from concept to completion.

Dr. Ishaan Gupta: Conceived and supervised the overall project; provided the original idea, supported funding in the dry lab experiment, and provided continuous guidance throughout all stages of the study, from concept to completion.

## Ethics Declaration

### Ethics approval and consent to participate

Not applicable.

## Consent for publication

Not applicable.

## Supplementary Information

Supplementary Document 1.: Bar chart shows the distribution of genomic regions based on their PhastCons conservation scores, with most regions falling into high-score bins, indicating strong evolutionary conservation across species.

Supplementary Document 2.: lnc-RNAs which are important for fin regeneration Supplementary Document 3.: lnc-RNAs which are important for fin positional memory Supplementary Document 4.: The list of overlapping lncRNAs

Supplementary Document 5.: A more detailed, integrative analysis is here where BAM files from various time points were merged with those from the positional memory dataset and normalized.

Supplementary Document 6.: WGCNA analysis was performed on this normalized matrix, generating a dendrogram with 89 color modules

Supplementary Document 7.: pathway enrichment analysis, and the enrichment maps (EMap) of selected module .

Supplementary Document 8.: The list of all regions and their corresponding modules

